# New hypotheses of cell type diversity and novelty from comparative single cell and nuclei transcriptomics in echinoderms

**DOI:** 10.1101/2022.05.06.490935

**Authors:** Anne Meyer, Carolyn Ku, William Hatleberg, Cheryl A. Telmer, Veronica Hinman

**Author notes:** Corresponding Author: Veronica Hinman.

## Abstract

Cell types are the fundamental building blocks of metazoan biodiversity and offer a powerful perspective for inferring evolutionary phenomena. With the development of single-cell transcriptomic techniques, new definitions of cell types are emerging. This allows a conceptual reassessment of traditional definitions of novel cell types and their evolution. Research in echinoderms, particularly sea star and sea urchin embryos have contributed significantly to understanding the evolution of novel cell types, in particular the primary mesenchyme cells (PMCs) and pigment cells that are found in sea urchin but not sea star embryos. This paper outlines the development of a gene expression atlas for the bat star, *Patiria miniata*, using single nuclear RNA sequencing (snRNA-seq) of embryonic stages. The atlas revealed 22 cell clusters covering all expected cell types from the endoderm, mesoderm and ectoderm germ layers. In particular, four distinct neural clusters, an immune cluster, and distinct right and left coelom clusters were revealed as distinct cell states. A comparison with *Strongylocentrotus purpuratus* embryo single cell transcriptomes was performed using 1:1 orthologs to anchor and then compare gene expression patterns. *S. purpuratus* primordial germ cell equivalents were not detected in *P. minata*, while the left coelom of *P. miniata* has no equivalent cell cluster in *S. purpuratus*. Pigment cells of *S. purpuratus* map to clusters containing immune mesenchyme and neural cells of *P. miniata*, while the PMCs of *S. purpuratus* are revealed as orthologous to the right coelom cluster of *P. miniata*. These results suggest a new interpretation of the evolution of these well-studied cell types and a reflection on the definition of novel cell types.

## Introduction

Cell types are the fundamental units of multicellular life. Understanding how novel cell types emerge is critical to understanding the relationships between cells, and hence the processes by which organisms acquire new evolutionary traits. Although fundamental to theories of morphological evolution, understanding how novel cell types evolve, or even how to define a novel cell type, is still unclear and a source of active debate (Arendt et al., 2016; Kin, 2015; Márquez-Zacarías et al., 2021). Traditionally a combination of morphology, function, cell lineage and subsets of gene expression have been used to describe and classify cell types. However, there are many ways in which this method of classification may not accurately reflect evolutionary relationships. For example, cell types with similar morphologies, such as the striated myocytes in bilaterians and cnidarians, appear to be homologous morphologically, when in fact this phenotype arose independently (Brunet et al., 2016). Developmental lineage too has drawbacks, as cell types that are very similar in regulatory profiles and morphology may arise from different cell lineages in an embryo. For example, neurons can derive from both the ectoderm and foregut endoderm in sea urchins (Angerer et al., 2011).

Gene regulatory networks (GRNs) are a valuable tool for understanding the evolution of cell types within and between species, as evolutionarily related cell types are assumed to have similar regulatory expression profiles prior to divergence by genetic individualization. Homologous cell types can therefore be identified between species by examining the expression of orthologous genes as a proxy for GRN operating in the cells. However, the cooption of modular GRN subcircuits may occur frequently during evolution, obscuring the relationship between cells in an organism (Monteiro and Podlaha, 2009) (Davidson and Erwin, 2010).

The phylum Echinodermata, which includes species of sea stars, sea urchins, brittle stars, sea cucumbers, and feather stars, are a powerful system for studying the evolution of cell types. Their larval forms in particular have several cell types that, based on developmental, morphological, and functional criteria, are considered to be novel (Figure 1). One of the best-studied examples of novel cell types in these species is the biomineral forming skeletal cells of sea urchin and brittle star pluteus larvae. These cells derive from early embryonic primary mesenchyme cells (PMCs) that ingress and secrete a calcium-based biomineral skeleton within the blastocoel. The long skeletal rods formed by this process give the larvae the distinctive pyramidal, plutei shape (Decker and Lennarz, 1988). The larval forms of the other groups of echinoderms—the bipinnaria in sea stars, auricularia in sea cucumbers, auricularia-like in crinoids, and the tornaria larva of the outgroup Hemichordate phylum—do not produce larval skeletal rods. The sea cucumber auricularia does however produce a small biomineralized cluster that appears to be secreted from mesenchyme cells (Decker and Lennarz, 1988; Hyman, 1955; McCauley et al., 2012).

**Figure 1:**
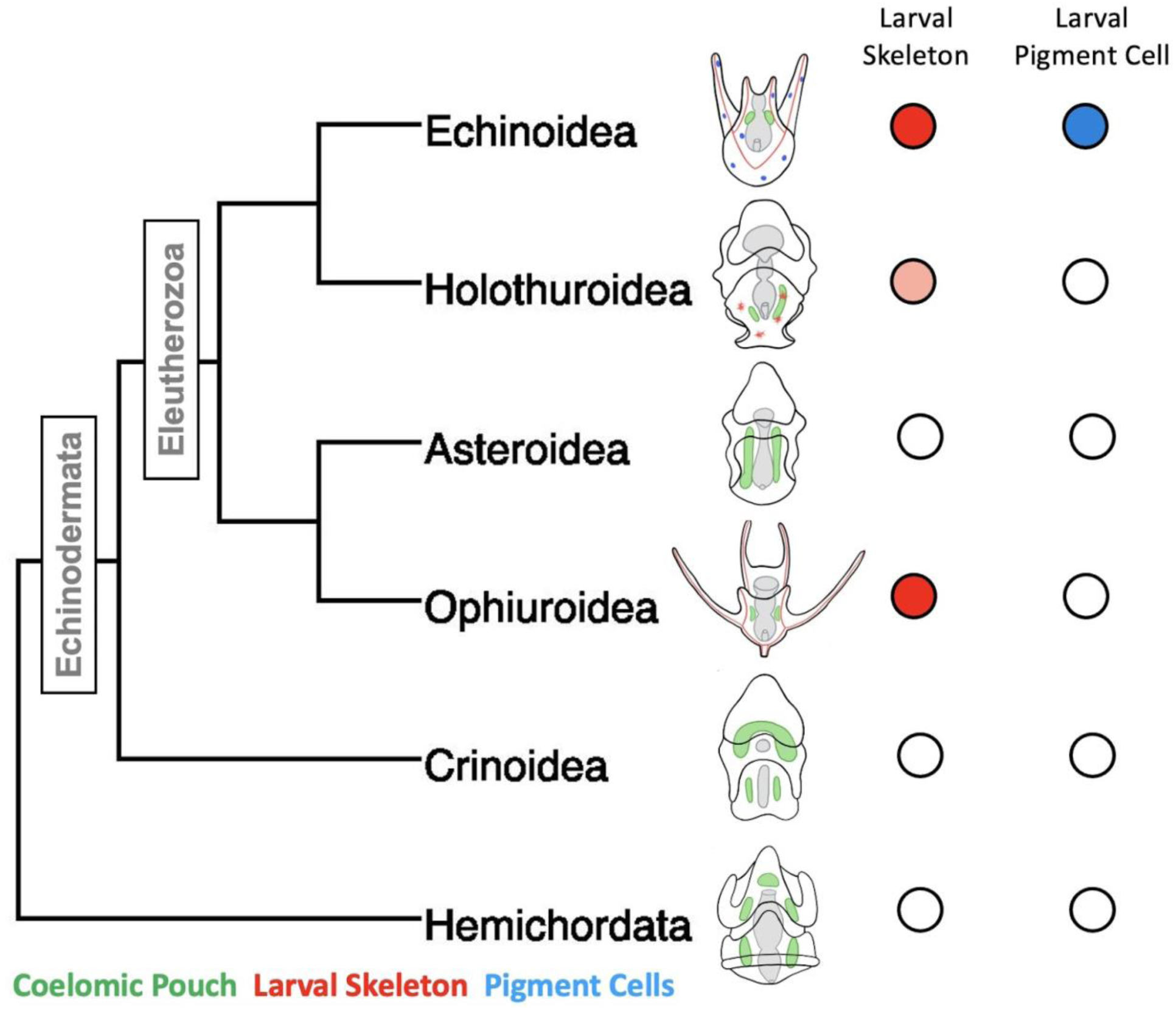
Within Echinodermata, two novel cell types have emerged. A phylogeny of the major classes of Echinodermata, and Hemichordata outgroup. Embryonic schematics show biomineralized larval skeletons colored in red, coelomic pouches colored green, and pigment cells colored in blue. Presence/absence is also indicated using colored circles to the left of the embryos. The pink shading indicates the secretion of biomineralized spicules, but not the formation of an extensive skeleton.

Another novel cell type, the larval pigment cells are unique to the sea urchins (i.e. the echinoid family). The pigment cells emerge in the second wave of mesenchymal ingression during gastrulation and imbed themselves in the ectoderm of the larvae (Gustafson and Wolpert, 1967; Massri et al., 2021). Upon the activation of an immune response, these cells ingress into the blastocoel and aggregate with other immune cells at sites of high pathogen concentration (reviewed in (Buckley and Rast, 2017). The molecule that gives these cells their distinctive color is echinocHrome A, a naphthoquinone with known antimicrobial properties (Service and Wardlaw, 1984).

Much is known about the mechanisms of specification of these novel cell types in sea urchins, especially the purple sea urchin, *Strongylocentrotus purpuratus,* and the green sea urchin, *Lytechinus variegatus*, which have an especially well-resolved early developmental GRNs.(Davidson et al., 2002a, 2002b; Hinman et al., 2003a; Oliveri et al., 2002; Saunders and McClay, 2014; Smith et al., 2018)). This GRN has also provided the basis for comparison with several other echinoderm taxa, most extensively the bat sea star, *Patiria miniata* (Cary et al., 2020; Hinman and Cheatle Jarvela, 2014; Hinman et al., 2009). Sea stars such as *P. miniata* morphologically lack both larval-skeleton-forming cells and pigment cells (Figure 1).

Comparisons of GRNs within different cell types have been used to infer the evolution of the sea urchin skeleton and pigment cell types from an ancestor shared with sea stars. A leading hypothesis from these studies is that the sea urchin and brittle star larval skeleton arose through the co-option of the adult skeletogenesis program by a subpopulation of embryonic mesodermal cells (Gao and Davidson, 2008; Morino et al., 2012), under the regulatory control of the transcription factor aristaless-like homeobox (*alx1*) and cooption of the vascular endothelial growth factor receptor (*VEGFR*) (Ben-Tabou de-Leon, 2022; Ettensohn and Adomako-Ankomah, 2019; Morino et al., 2012). These regulatory genes are required for the expression of genes needed for biomineralization in these sea urchin cells (Duloquin et al., 2007; Rafiq et al., 2012). *VEGFR*in particular has not been detected in echinoderm embryos that do not form skeletons. Likewise, the pigment cells are also proposed to have also arisen through a heterochronic co-option of an adult program for the formation of these cells (Perillo et al., 2020), although the possible evolution of this cell type is still poorly investigated.

Single-cell RNA-sequencing (scRNA-seq) has revolutionized our understanding of cell-type diversity within an organism (Konstantinides et al., 2018; Marioni and Arendt, 2017; Morris, 2019; Shapiro et al., 2013; Tanay and Sebé-Pedrós, 2021). This technology profiles the transcriptome of individual cells within an organism, preserving the heterogeneity of cell states that are lost in traditional bulk sequencing. This new technology provides prominence to gene expression and gene regulatory networks (GRNs) as the basis of cell-type identity and the evolutionary relationships between cell types (Achim and Arendt, 2014; Arendt, 2008; Arendt et al., 2016). Knowledge of echinoderm cell diversity and molecular profiles has recently benefited greatly from scRNA-seq studies in echinoids (Foster et al., 2020; Massri et al., 2021; Paganos et al., 2021). These datasets provide an unbiased view of the gene expression state of cell types in sea urchin embryos. This presents an opportunity to establish equivalent profiling in sea stars to establish the similarities and differences of gene expression states on a large scale rather than the candidate gene approaches that have been performed to date. We, therefore, developed an snRNA-seq developmental cell atlas for *P. miniata*. Single-nucleus RNA-seq (snRNA-seq), is performed on isolated nuclei rather than whole cells (Lake et al., 2016) and is suggested by recent literature to provide a more representative view of cell types than single-cell sequencing because the uniform nuclear morphology permits less biased isolation (Wu et al., 2019). This analysis for the first time provides a global view of cell state expression profiles in sea stars.

To compare expression states across species, we made a multi-species developmental atlas from previously published *S. purpuratus* scRNA-seq data (Foster et al., 2020) using an integrated approach based on the expression patterns of a comprehensive mapping of 1:1 gene orthologs between *S. purpuratus* and *P. miniata* (Beatman et al., 2021; Foley et al., 2021). For each gene discussed in this paper, the species’ LOCIDs and associated nomenclature based on (Beatman et al., 2021) are presented in supplemental table 1. We use these atlases to compare the GRN definition of cell type novelty to the morphological definition to establish alternative hypotheses of the origin of pigment cells and PMCs and to broadly reflect on the definitions of cell type and novelty.

## Results

### Single-nucleus transcriptome is representative of cell diversity in sea star embryos

Before creating our *P. miniata* atlas, we wanted to explore the use of snRNA-seq in echinoderms to see if it offers any benefit over scRNA-seq, including more robust isolation. We, therefore, adapted protocols for single nuclear isolation used in Omni-ATACseq (Corces et al., 2017). This is the first time that a single nuclei sequencing protocol has been conducted in an echinoderm.

We sought to establish whether our nuclear isolation method was less biased in cell-typer etention by comparing bulk-RNA seq data collected from samples isolated using whole-cell isolation, nuclear isolation, or from whole embryos (Figure 2.A). We compared the overall similarity of transcriptome profiles using the Spearman correlation coefficient for each dataset (Figure 2.B). The nuclei to whole embryo comparison had the highest Spearman coefficient (⍴= 0.987), indicating higher overall transcriptome similarity to the undissociated embryos. Whole-cell isolation also showed a strong correlation with the whole embryo, with a spearman coefficient of ⍴=0.9857839, demonstrating that, like nuclear isolation, single-cell isolation still reflects the overall transcriptomic profile of the whole organism.

**Figure 2:**
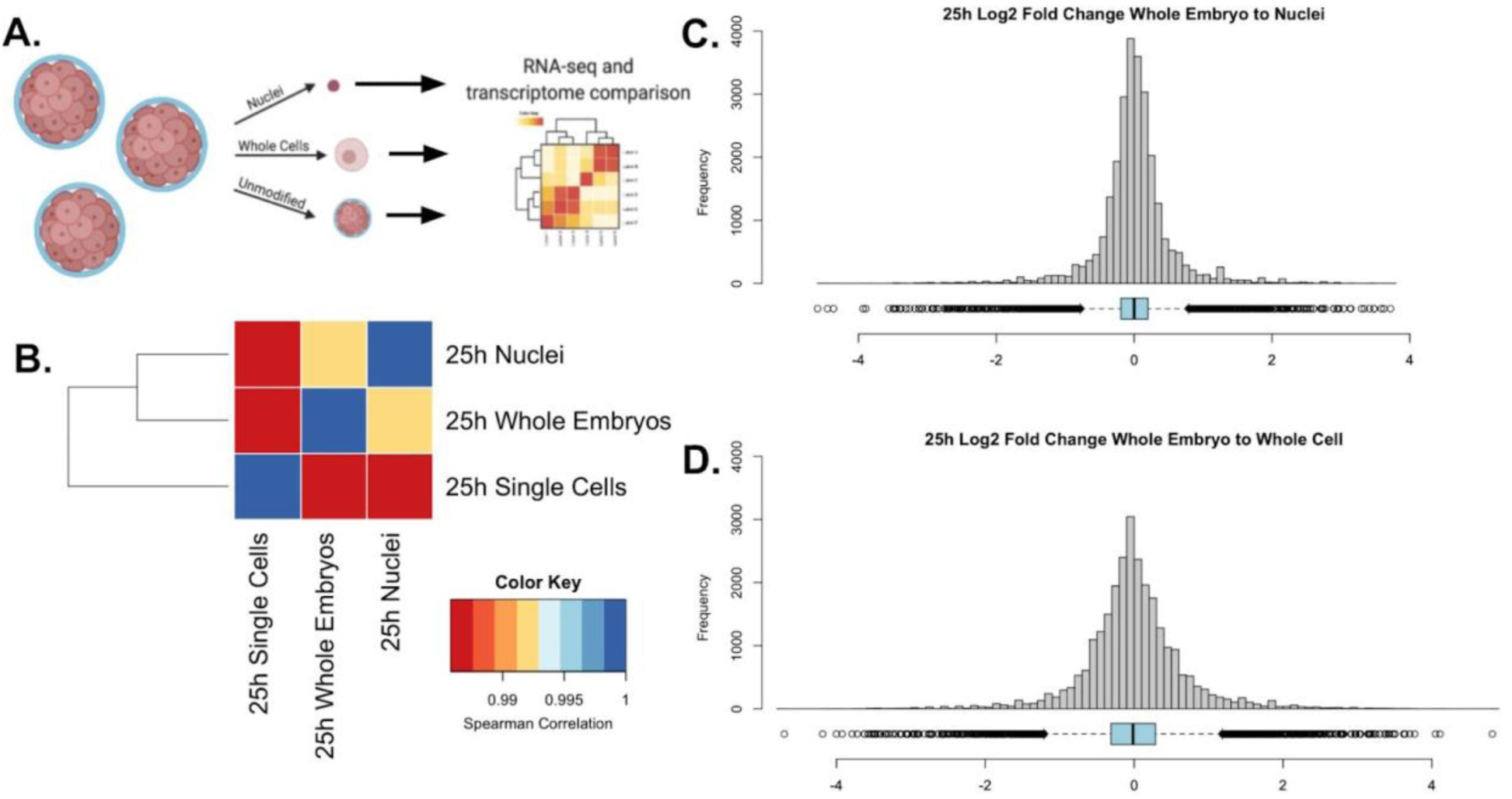
Nuclear isolation more robustly captures cell diversity than whole-cell extraction methods. (A) A schematic representation of the experimental design. A single culture of 25 hpf *P. miniata* embryos was partitioned into three aliquots: one left unmodified, one underwent nuclear extraction, and the other underwent whole-cell extraction. Bulk RNA-seq was performed on each sample and their transcriptomes were compared across samples. (B) Spearman correlation was conducted on quantile-normalized gene counts from the RNA-seq dataset. Nuclear isolation had the highest correlation with the whole embryo RNA-seq dataset with ⍴=0.9916922. The whole-embryo to the whole-cell spearman correlation coefficient of ⍴=0.9857839. Whole-cells to nuclei has a coefficient of ⍴= 0.9867828. (C & D) Histograms and dot plots display the frequency distribution of log2 fold changes in gene expression levels between the whole embryo and nuclear or whole-cell data sets. (C)The nuclear RNA-seq datasets resulted in a distribution where *μ*=0.013, *σ*=0.558, and M=0.000.(D) The whole-cell to whole embryo comparison resulted in a distribution where *μ*=-0.003, *σ*=0.683, and M=-0.013.

To further assess nuclear vs. cellular isolation bias, we calculated the log2 fold change in gene transcript count for each of our isolation methods with respect to the intact embryo control (Figure 2.C-D). The nuclear to whole-embryo comparison resulted in a mean of 0.013 and a standard deviation of 0.558. The whole-cell to whole-embryo comparison resulted in a mean of - 0.003 and a standard deviation of 0.683. Though the mean is closer to zero in the whole-cell isolates, we conclude that a nuclear isolation method still more consistently reflects whole embryo expression diversity because of the lower standard deviation, indicating the overall distribution is more narrow and therefore more genes accurately reflect the expression levels of a whole organism.

Finally, as proof of principle, we performed snRNA-seq in *S. purpuratus* to determine if we could identify distinct clusters of known cell types. Using samples from four time points (6 hours post-fertilization (hpf), 15 hpf, 23 hpf, and 33 hpf), we were able to identify 15 clusters, encompassing all four germ layers and including several clusters corresponding to PMCs and pigment cells. These PMC clusters were characterized by gene expression associated with different degrees of PMC differentiation. Further analysis of this pilot dataset is provided in Supplementary Text Section 1 and Supplementary Figure 1).

### A cell type atlas for *Patiria miniata* embryo development

Previous work has shown that pooling scRNA-seq data from the 8-cell stage to the late gastrula of *S. purpuratus* into one atlas results in biologically meaningful clustering (Foster et al., 2020). Therefore, we conducted snRNA-seq at three different time points within an early developmental window spanning from blastula to mid gastrula stage, to compile an atlas describing early development in *P. miniata.* snRNA-seq was conducted on *P. miniata* embryos at three different time points (16hpf, 26hpf, 40hpf) (Figure 3.B). These timepoints span from blastula to mid gastrulation when cell types of the larva are first specified (Cary et al., 2020; Yankura et al., 2010).

**Figure 3:**
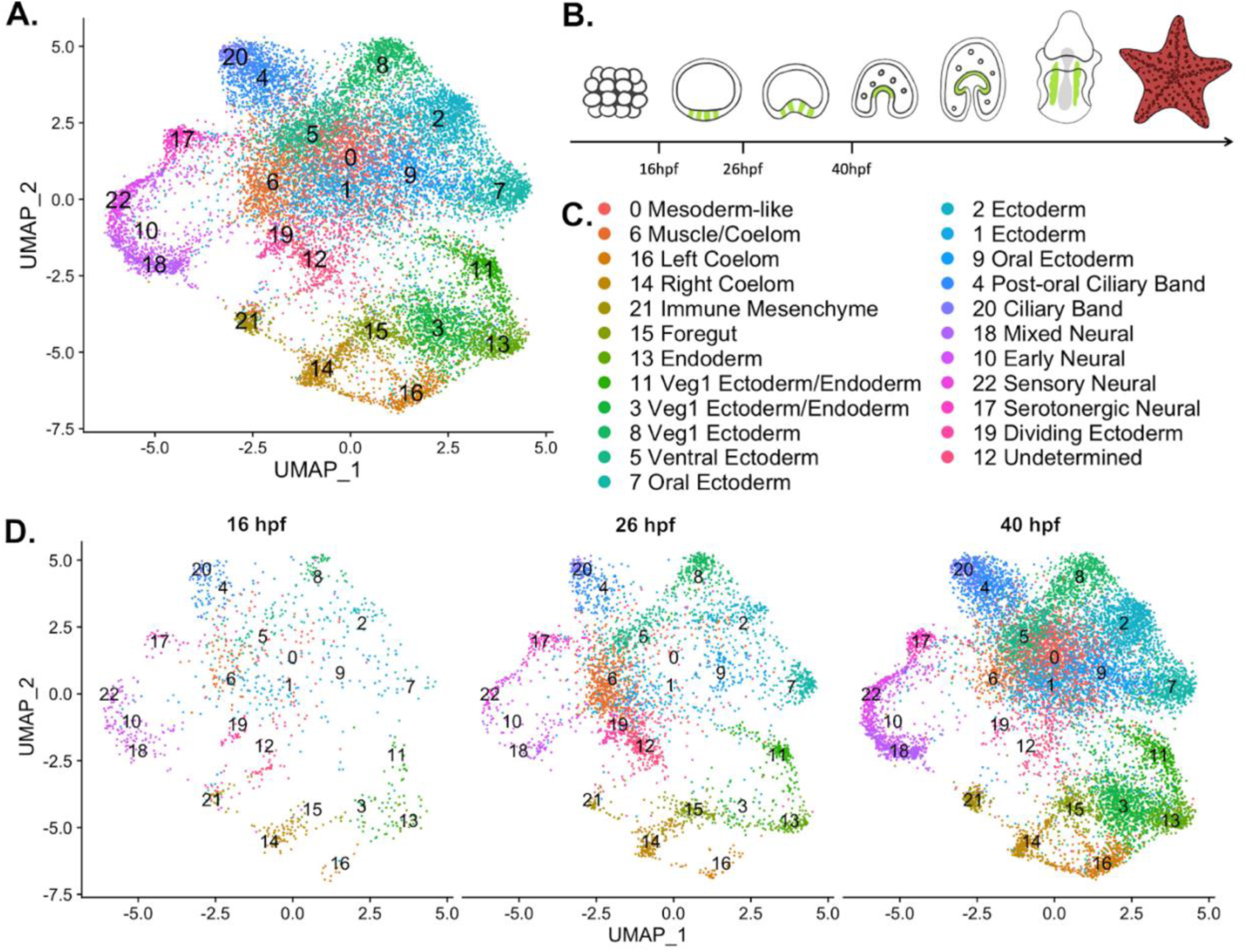
A nuclear atlas for early *P. miniata* development. (A) The UMAP projection of the *P. miniata* atlas developed from snRNA-seq of 3 developmental time points. 17,118 nuclei were considered in total. (B) The schematic shows time points sampled. The green color indicates the coelomic pouches. (C) A key displaying the cluster names corresponding to UMAP projections Figure 3.A and Figure 3.D. (D) The UMAP projection of the *P. miniata* nuclear atlas, separated by the 3-time points of sampling, 16hpf has 1076 nuclei, 26hpf has 3888 nuclei, and 40 has 12203 nuclei.

We isolated nuclei and performed single nuclear sequencing, processed reads, and aligned them to a pre-mRNA index of the recently assembled *P. miniata* genome v3.0 (Arshinoff et al., 2022). After data filtration and quality control (see Methods), we were left with 17,188 nuclei. Dimensional reduction and clustering (see Methods) resulted in 23 clusters (abbreviated with Pmin CL) (Figure 2.A). Cluster size ranged from 133 nuclei (Pmin CL 22 “Sensory Neural”) to 1739 nuclei (Pmin CL 0 “Mesoderm-like”).

All clusters contain nuclei from each of the time points indicating that there are no substantial batch effects (Figure 3.D and Supplementary Figure 2). However, there are some time point enrichments found in the clusters. Clusters 7, 18, and 22 are dominated by cells from the later time points, with only 10.6%, 3.27%, and 5.09% cluster members originating from the earliest time point respectively. Clusters 10, 14, and 21 are dominated by cells from the earlier time points, with 63.0%, 49.6%, and 56.1% originating from the 16 hpf dataset. These observations suggest that differences in developmental timing may have an impact on transcriptome profiles and therefore should be kept in mind during cluster annotation.

Representatives from all germ layers were identified through cluster annotation, as is displayed in Figure 4. This figure presents a subset of genes, whose expression patterns were used to inform the naming of each cluster. Cluster annotation was performed by calculating genes that are differentially expressed in a cluster relative to the rest of the dataset, henceforth known as marker genes (see methods for marker gene calculation and classification). Additionally, genes with known expression patterns (from previously published literature) and highly conserved cell-type-specific functions (such as enzymes involved in neurotransmitter synthesis) also helped inform cluster annotation. Finally, we performed whole-mount in situ hybridization (WMISH, see methods) on a select group of genes to validate cluster annotations (Figure 5-7).

**Figure 4:**
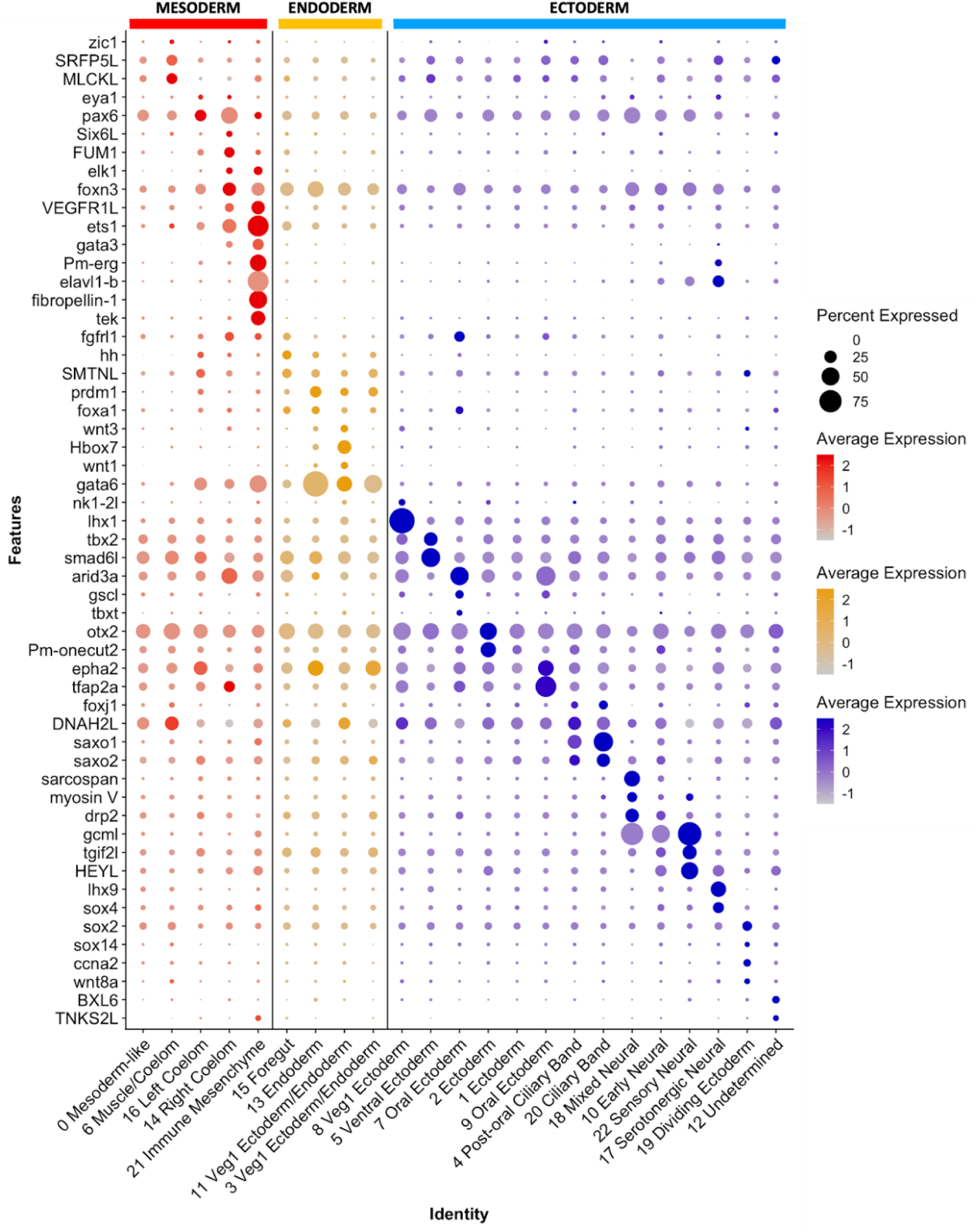
Differential expression of marker genes was used to annotate cluster identity. A dot plot highlighting marker genes used to determine cluster identity. The X-axis lists the cluster names, while the Y lists gene names. Mesodermal cells are colored in red, endoderm in yellow, and ectoderm/undetermined in blue. Circle size corresponds to the number of cells in the cluster expressing the gene of interest, while shade correlates with the level of expression.

**Figure 5:**
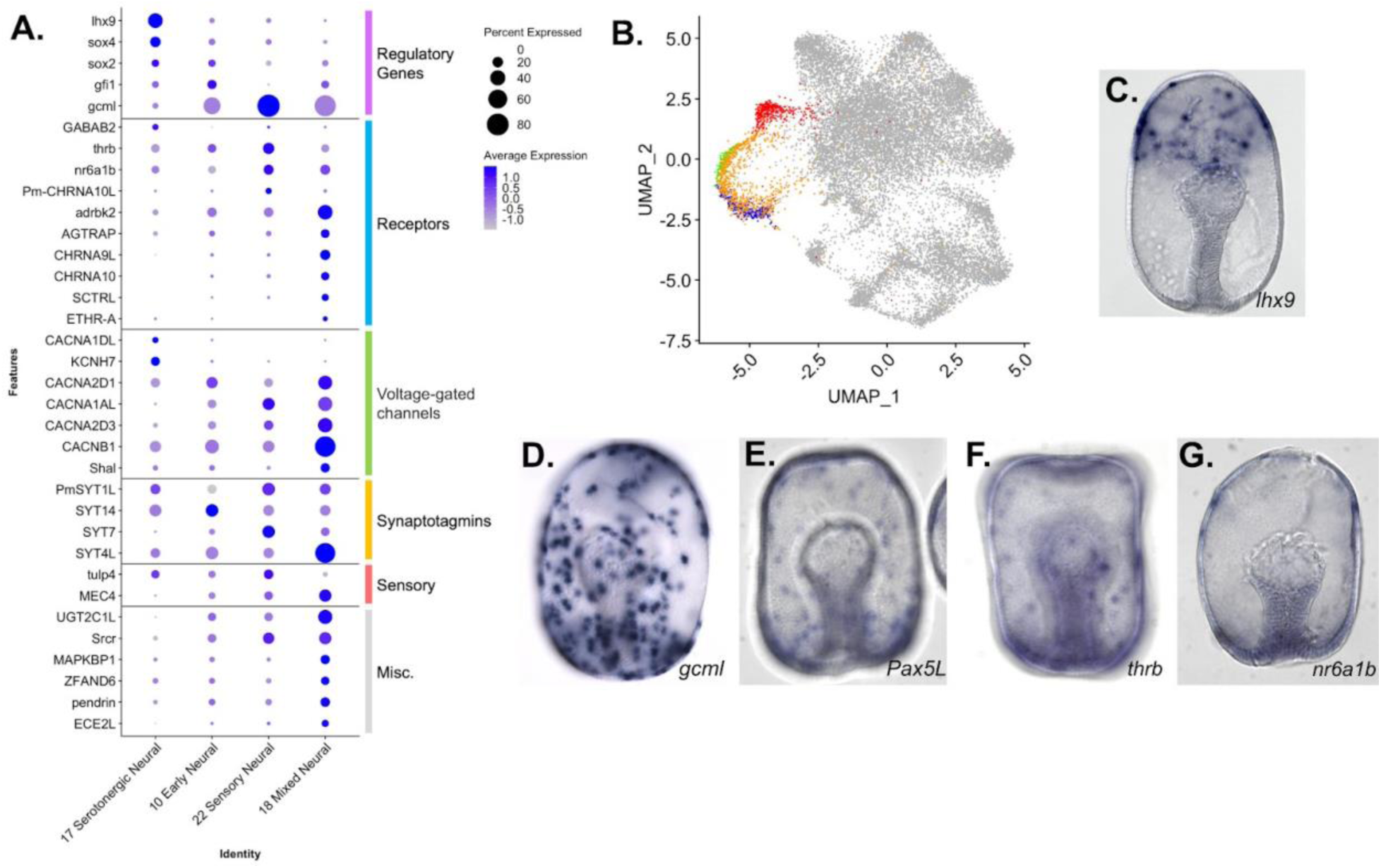
Characterization of neuronal populations of the *P. miniata* Atlas. (A) A dot plot of the four neural clusters and a selection of genes used to annotate the clusters. The genes are divided into six categories: regulatory genes, receptors, voltage-gated channels, synaptotagmins, sensory, and misc. (B) A UMAP projection highlights the four neural clusters. (C-H) WMISH validation of marker gene expression at the gastrula stage. (C) *Lhx9* is expressed in serotonergic neuronal precursors near the animal pole. (D) *gcml* is expressed in cells throughout the ectoderm, with an exclusion zone on the oral, non-neurogenic region of the embryo. (E) *Pax5L* is expressed in cells embedded throughout the ectoderm, with an exclusion zone on the non-neurogenic oral surface (observed 46/46 embryos) (F). *Thrb* is expressed in cells embedded throughout the ectoderm(observed 32 /33). (G) *nr6a1b* is expressed in cells embedded in the ectoderm (observed 38/38).

Seven of the 23 identified clusters (Pmin CL ID 17, 22, 18, 10, 21, 14, and 16), are discussed in detail below and will be referred to in our subsequent comparative analysis. Four of these clusters (Pmin CL 17, 22, 18, 10) represent neuronal cell types, offering valuable insight into the diversity of neurons in early echinoderm embryonic development. One represents a distinct immune cell cluster (Pmin CL 21), which is the first time this cell type has been molecularly characterized in this species. Finally, we identified two populations of coelomic cells (Pmin CL 14 and 16), indicating that these tissues are composed of transcriptomically distinct populations. A complete description of the cluster annotation justification for clusters not discussed in this paper’s body is provided in Supplementary Text Section 2.

### Neural clusters

The four neural clusters (Pmin CL 17, 22, 18, 10) are all closely associated in the UMAP projection (Figure 5.A). Their identities were established based on the expression of known neural markers in *P. miniata* and genes with well documented, conserved functionality in neural function including pre-synaptic synaptotagmins, neurotransmitter receptors, and the presence of voltage-gated ion channels (Adolfsen and Littleton, 2001; Burke et al., 2006; Moghadam and Jackson, 2013; Walker and Holden-Dye, 1991) (Figure 5.B).

The marker gene sets for these four neural clusters include four synaptotagmin genes: synaptotagmin-7 (*SYT7*), synaptotagmin-14 (*SYT14*), synaptotagmin-1-like (*PmSYT1L*), and synaptotagmin-like protein 4 (*SYT4L*). The synaptotagmin gene family has been extensively documented to have conserved functionality in neurons across phyla (Adolfsen and Littleton, 2001). *SYT7* is most highly expressed in Pmin CL 22. *SYT14* is expressed in all neural clusters but is most highly expressed in Pmin CL 10. Conversely, *SYT1L* is expressed in all clusters but shows the lowest expression in Pmin CL 10. S*YT4L* is expressed in all clusters, but most strongly in Pmin CL 18 (Figure 5.A). This suggests that in echinoderms, members of the synaptotagmin gene family are differentially expressed between different classes of neurons, as they are in Drosophila and Mammalia (Adolfsen et al., 2004; Craxton, 2004; Marquèze et al., 1995).

### Serotonergic Neural

LIM homeobox 9 (*lhx9)*, a gene previously established as a marker of serotonergic neurons (Cheatle Jarvela et al., 2016), was found to be a strong marker of Pmin CL 17 (adjusted p-value = 5.45E-190) and was verified using WMISH (Figure 5C). Tryptophan 5-hydroxylase 1-like (*TPH1L*), a gene involved in serotonin biosynthesis (Fitzpatrick, 1999) has been shown to localize to serotonergic neurons in sea urchins, (Yaguchi and Katow, 2003), was also identified as a marker of this cluster (adjusted p-value = 1.28E-09). This along with the expression of other known markers including transcription factor SRY-box transcription factor 4 (*sox4*, adjusted p-value = 2.24E-29), neogenin-like (*NEO1L*), and ELAV-like protein 1B (*elavl1-b*, adjusted p-value = 2.50E-72) allow us to conclude Pmin CL 17 corresponds to serotonergic neurons.

### Sensory Neural

Pmin CL 22 was annotated as a sensory neuron population. Genes relating to neurotransmitter receptors were also found to be expressed in this cluster. These include neuronal acetylcholine receptor subunit alpha-10-like (*CHRNA10L*, adjusted p-value = 3.14E-15), and beta-adrenergic receptor kinase 2 (*adrbk2*, adjusted p-value = 1). It was also marked by the expression of thyroid hormone receptor beta (*thrb*, adjusted p-value = 0.17) and nuclear receptor subfamily 6 group A member 1B (*nr6a1b*, previously known as steroid hormone receptor 3, adjusted p-value = 1). WMISH of these two genes confirmed localization to a subset of ectoderm, consistent with a neural population (Figure 5.F-G). This cluster was also represented by high levels of the transcription factor Paired box protein Pax5-like (*Pax5L*, adjusted p-value =4.07E-299), which was also shown via WMISH to express in cells embedded in the ectoderm (Figure 5.E).

Finally, Pmin CL 22 was marked by genes related to a diverse range of sensory capabilities such as mechanosensing (degenerin mec-4 (*MEC4*, adjusted p-value = 3.06E-19) (O’Hagan et al., 2005) and photosensing (TUB like protein 4 (*tulp4*, adjusted p-value =0.01641617)) (Carrella et al., 2020).

Glial cells missing transcription factor-like (*gcml*), the sea star ortholog of the Drosophila glial cells missing (Dmel_CG12245), was also identified as a statistically significant marker gene of Pmin CL 22 (adjusted p-value = 1.04E-250). WMISH verified that this gene localizes to cells embedded in the ectoderm (Figure 5.D).

### Mixed Neural

Pmin CL 18 corresponded to a population of cells of mixed neural specification. As with Pmin CL 22, this population is distinct in its higher expression of *gcml* and *Pax5L* (adjusted p-value= 1.62E-199). It also is defined by the expression of neurotransmitter receptors neuronal acetylcholine receptor subunit alpha-9-like (*CHRNA9L*, adjusted p-value = 2.51E-200), neuronal acetylcholine receptor subunit alpha-10 (*CHRNA10*, adjusted p-value = 2.90E-146) and hormonal receptors including *nr6a1b* (adjusted p-value = 9.43E-05), type-1 angiotensin II receptor-associated protein-like (*AGTRAP*, adjusted p-value = 4.44E-19) and ecdysis triggering hormone receptor subtype-A (*ETHR-A*, adjusted p-value = 7.86E-19). Several other genes related to neuroendocrine function were also present, such as endothelin-converting enzyme 2- like (*ECE2L*, adjusted p-value = 5.22E-96) (Mzhavia et al., 2003), and pendrin(adjusted p-value = 1.49E-14) (Royaux et al., 2000).

Pmin CL 18 is also marked by the expression of serotonin N-acetyltransferase-like (AANAT, adjusted p-value = 2.46E-89). Pmin CL18 is also characterized by the expression of highly conserved muscle-associated genes such as dystrophin-related protein 2 (*drp2*, adjusted p-value = 3.66E-33) (Huang et al., 2004) and sarcospan (adjusted p-value = 1.25E-212) (Hooper and Thuma, 2005; Lehman and Szent-Györgyi, 1975).

Genes potentially related to immune function were also detected as markers including UDP-glucuronosyltransferase 2C1-like (*UGT2C1L*, adjusted p-value =4.79E-67) (Wang et al., 2021), Scavenger receptor cysteine-rich domain superfamily protein (*Srcr*, adjusted p-value = 2.52E-43) (Pancer et al., 1999) and several genes linked to the NFκB pathway (mitogen-activated protein kinase-binding protein 1 (*MAPKBP1*, adjusted p-value = 1.68E-15)) (Fu et al., 2015) and AN1-type zinc finger protein 6 (*ZFAND6*, adjusted p-value = 7.23E-10)) (Huang et al., 2004).

### Early Neural

Finally, Pmin CL 10 was also annotated as a non-specific early neuronal population. This cluster is marked by the expression of *Pax5L* (adjusted p-value = 1.12E-145) and beta-adrenergic receptor kinase 2 (*adrbk2*, adjusted p-value = 6.82E-16). Though not a marker gene, this cluster also expresses *AGTRAP*. When we examine the normalized contribution of the three sampling time points to each cluster (Supplementary Figure 2), we see that our earliest time point (6 hpf) makes up 63% of cluster 10, as compared to 5% of cluster 22, 3% of cluster 18, and 34% of cluster 17. Therefore, Pmin CL 10 may represent an earlier population of neurons, whereas the other three neural clusters are more specified neuronal populations.

### Immune Mesenchyme

Pmin CL 21 is a very spatially distinct cluster in the UMAP dimensional reduction plot (Figure 5.A). We annotated this cluster as mesenchymal in nature based on the expression of known mesenchymal markers, i.e. ETS proto-oncogene 1 transcription factor (*ets1*), transcriptional regulator ERG homolog (*Pm-erg*), and *elavl1-b* (Cheatle Jarvela et al., 2016; Hinman and Davidson, 2007; McCauley et al., 2010). We confirmed that these genes are expressed in mesenchymal populations using WMISH (Figure 6.D). We also identified Gata binding protein 3 (*gata3*) as a marker of this cluster (adjusted p-value =1.45E-211). Previous work in sea urchins has shown *gata3* plays a key role in blastocoelar immunocyte migration and maturation (Pancer et al., 1999). The WMISH presented in Figure 6.F confirms that this gene is similarly expressed in patches of the mesodermal archenteron, consistent with marking pre-mesenchymal cells.

**Figure 6:**
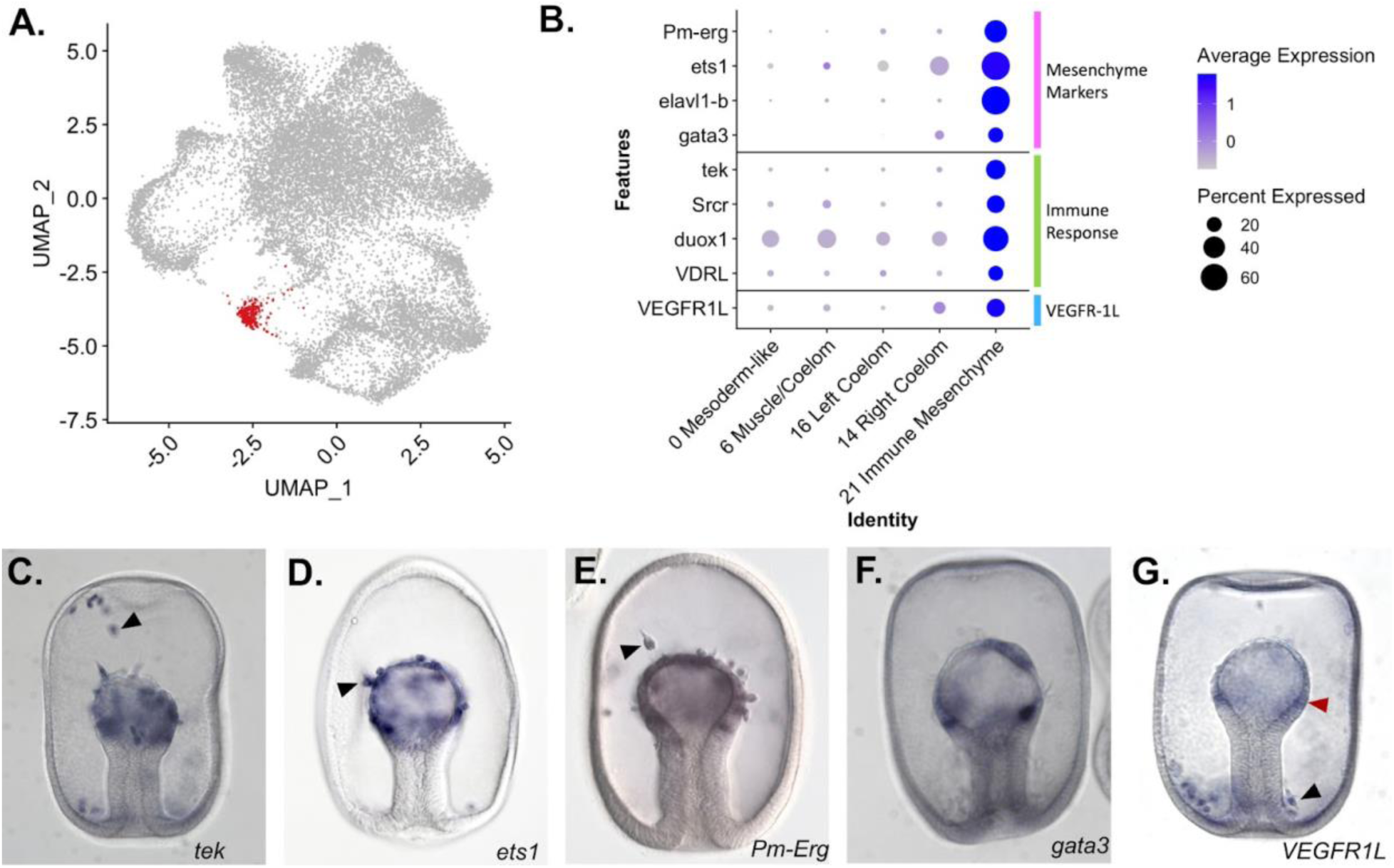
Identification of an Immune Mesenchyme cell type in *P. miniata*. (A) A UMAP projection highlights in red the cluster we annotate as immune mesenchyme. (B) A dot plot of mesodermal clusters and the genes used to identify the immune mesenchyme cluster identity. (C-G) WMISH validation of marker gene expression at the gastrula stage. (C) *tek* is expressed in the mesenchyme (indicated with arrow) and epithelial mesoderm (observed 33/34). (D) *Ets1* is expressed in mesodermal cells undergoing the epithelial to mesenchyme transition (indicated with arrow). (E) *Pm-Erg* is expressed in mesenchymal cells (indicated with arrow) as well as in the epithelium of the archenteron. (F) *Gata3* is expressed in patches of the archenteron epithelium (observed 27/ 30). *VEGFR1L* is expressed in the mesenchyme (indicated with black arrow) and archenteron epithelium (indicated with red arrow) (observed 51/55). Additional in situs showed that the VEGFR1L+ mesenchyme clustered in a ring around the base of the archenteron (observed in 28/36).

**Figure 7:**
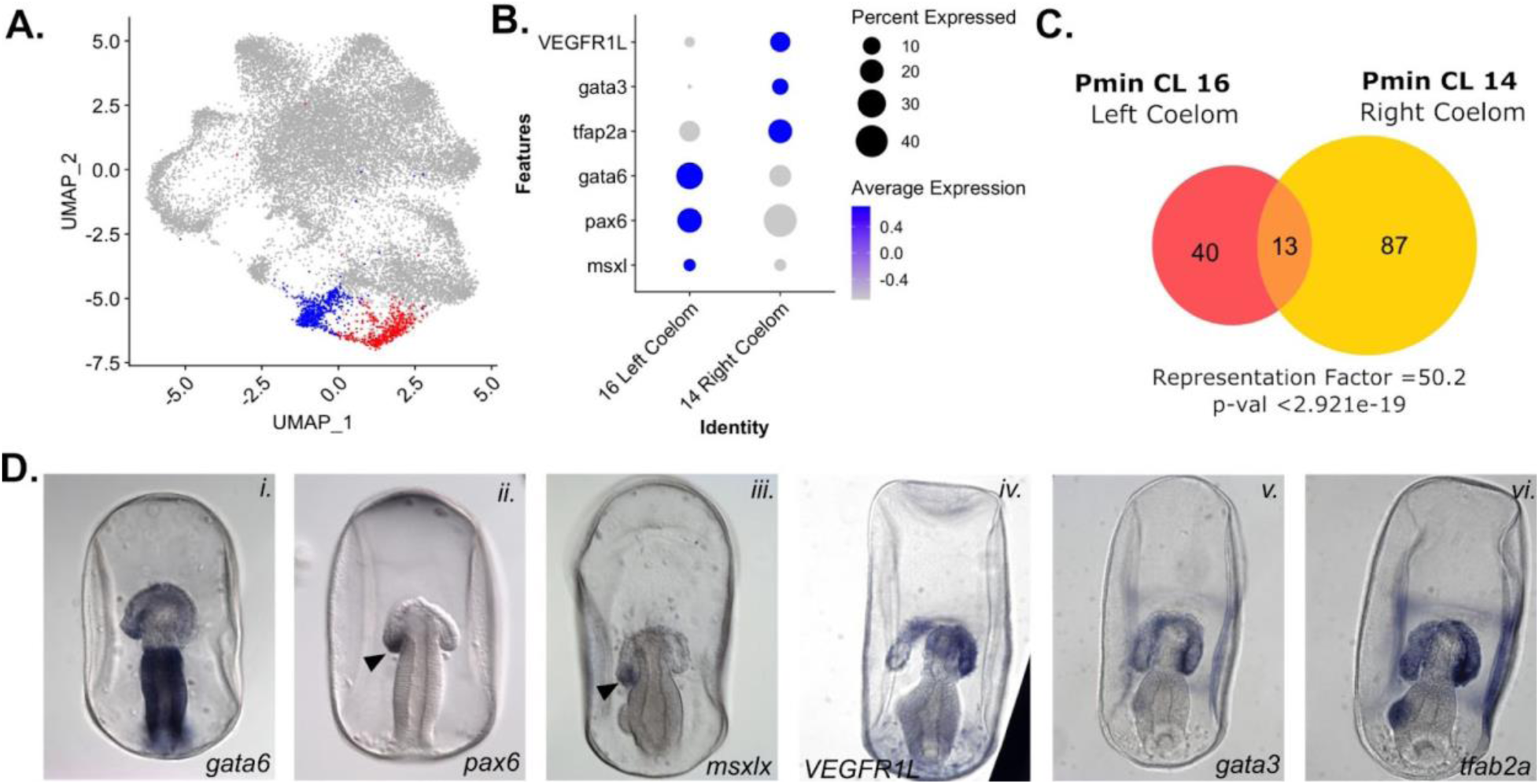
snRNA-seq detects left/right asymmetry in coelom development. (A) A UMAP projection of the *P. miniata* atlas highlights in blue the right coelomic pouch and in red the left colonic pouch. (B) A dot plot of two coelom clusters and the genes identified as differentially expressed between the two clusters. (C) A Venn diagram shows the overlap in marker genes shared between the two coelomic cells. 13 genes are shared in total, with the overlap yielding a representation factor of 50.2 with a p-value < 29.21e-19. (D) Validation of genes marking the left (i. - iii.) and right (iv. - vi.) coelomic clusters using WMISH in the late gastrula stage. *gata3, pax6*, and *msxlx* are expressed in the left coelomic pouch (indicated with arrow in ii. and iii.) and absent in the right pouch. *VEGFR1L*, *gata3*, and *tfap2a* are expressed in both pouches but are more highly expressed in the right coelomic pouch compared to the left.

The nuclei of this cluster also express genes involved in immune response in other echinoderms. TEK receptor tyrosine kinase (*tek*) was identified as a statistically significant marker (adjusted p-value = 1.65E-213). The ortholog of *tek* in *S. purpuratus* is expressed in coelomocytes and is upregulated in the apical organ during an immune challenge in adults (Stevens et al., 2010), indicating an immune function in sea urchins. *Srcr* (adjusted p-value = 6.46E-43) is also expressed in this cluster. A homolog of this gene in the asteroid *Asteria pectinifera* has been directly linked to larval immune response (Furukawa et al., 2012).

Genes linked to immune function in other animals, including dual oxidase maturation factor 1 (*duox1*, adjusted p-value = 1.52E-53) (Rada and Leto, 2008), ras-related protein Rap-2c (*Rap2*c, adjusted p-value = 0.00011995) (Gillespie et al., 2022), and Vitamin D3 Receptor-like (*VDR*L, adjusted p-value = 2.87E-20) (Newmark et al., 2017), were also detected as markers of this cluster.

We also identified vascular endothelial growth factor receptor 1-like (*VEGFR1L*) as a marker of Pmin CL 21 (adjusted p-value = 7.85E-57). WMISH of this gene (Figure 6.G) shows it localized to the mesodermal cells at the top of the archenteron and in a mesenchymal population that accumulates around the base of the archenteron, similar to the localization of PMCs in *S. purpuratus* (Duloquin et al., 2007).

### Coelomic clusters

Two clusters, Pmin CL 14 and 16, were annotated as coelomic mesoderm (Figure 7.A). *P. miniata* larvae have large coelomic pouches (Figure 1). These are formed from the mesoderm at the blastula stage, which buds off from the top of the archenteron during late gastrulation, and finally diverges into morphologically distinct left and right pouches. The left coelomic pouch will form the adult water-vascular system after metamorphosis (Child, 1941).

To examine the similarity between their marker gene sets, we calculated the representation factor, a measure of the significance of the overlap between two sets of objects, compared to what would be expected by chance. Values greater than 1 indicate greater than expected overlap between two sets, while values less than 1 indicate less than expected overlap.

These two clusters share 13 marker genes in common, with a representation factor value of 50.2 and p-val < 2921e-19, representing a high degree of similarity. Many of those shared genes are related to the extracellular matrix or cell/cell adhesion, such as collagen alpha-2(IV) chain-like (*col4a2l*), *fibulin-1*, collagen type IV alpha 1 chain (*col4a1*), mucin-16-like (*MUC16*), teneurin-3-like (*TENM3L*), and signal peptide, CUB and EGF-like domain-containing protein 1 (*SCUBE1*). Curiously, there are few regulatory genes shared by these populations at this time point, with only transcription factor AP4 (*tfap4*) and forkhead box p4 (*foxp4*) in common. These overlaps suggest that both coelomic pouches share similarities in tissue structure.

Pmin CL 16 is distinguished from Pmin CL 14 by the differential expression levels of paired box 6 (*pax6*), GATA binding protein 6 (*gata6*) and msh homeobox-like (*Msxl*) (Figure 7.B). *Gata6* (Figure 7.D.i) is expressed more broadly in the left coelomic pouch but is absent from the tip of the right coelomic pouch. WMISH of these genes in later gastrula stages shows that *pax6* (Figure 7.D.ii) and *Msxl* (Figure 7.D.iii) are at the tip of the left coelomic pouch and absent from the right. Therefore we annotated Pmin CL 16 as the Left Coelom.

*gata3, VEGFR1L,* and *ets1,* which were markers of the immune mesenchyme, Pmin CL 21 (Figure 5.B), were also found to be highly expressed in Pmin CL 14, relative to Pmin CL 16. WMISH revealed strong localization of *VEGFR1L* and *gata3* to the right coelomic pouch (Figure 7.D.vi-v) in ad. AT-rich interactive domain-containing protein 3A (*arid3a*, formerly referred to as protein dead ringer homolog) was also identified as a marker (adjusted p-value = 2.28E-23) of this right coelomic region. *arid3a* has been shown to be involved in the empithelial to meenchye transition (EMT) and primary mesenchymal formation in other echinoderms (Amore et al., 2003). The expression of these genes associated with mesenchymal transitions suggests that the right coelomic cluster is the point of origin of mesenchymal cells.

Although our time points of sampling pre-date recognizable left/right asymmetry in the *P. miniata* embryo, we still clearly see transcriptomic divergences that become morphologically observable later in development.

### *S. purpuratus* single-cell atlas from equivalent stages

In order to compare *P. miniata* to *S. purpuratus,* cell cluster identities an integrated atlas was necessary. An extensive comparison of whole embryo RNA-seq time series data between *S. purpuratus* and *P. miniata* has shown that *P. miniata* develops at a faster rate and that *S. purpuratus* (hatched and mesenchymal blastula stages) are equivalent in developmental timing to the *P.miniata* time points used to generate the atlas here (Gildor et al., 2019). We, therefore, first constructed an *S. purpuratus* atlas using the hatched and mesenchymal blastula stages from a previously published dataset (Foster et al., 2020). Our pilot snRNA-seq data was not used for the multi-atlas construction, as it was only a small exploratory data set. After quality control, 7,370 cells were selected for analysis and clustering. UMAP dimensional reduction and Louvain community assignment resulted in 20 distinct clusters, assigned the prefix Spur CL (Figure 8.A). Clusters were annotated following the same marker gene method used for *P. miniata* (see methods) and a dot plot of a subset of our most useful markers is presented in Supplemental Figure 3.

**Figure 8:**
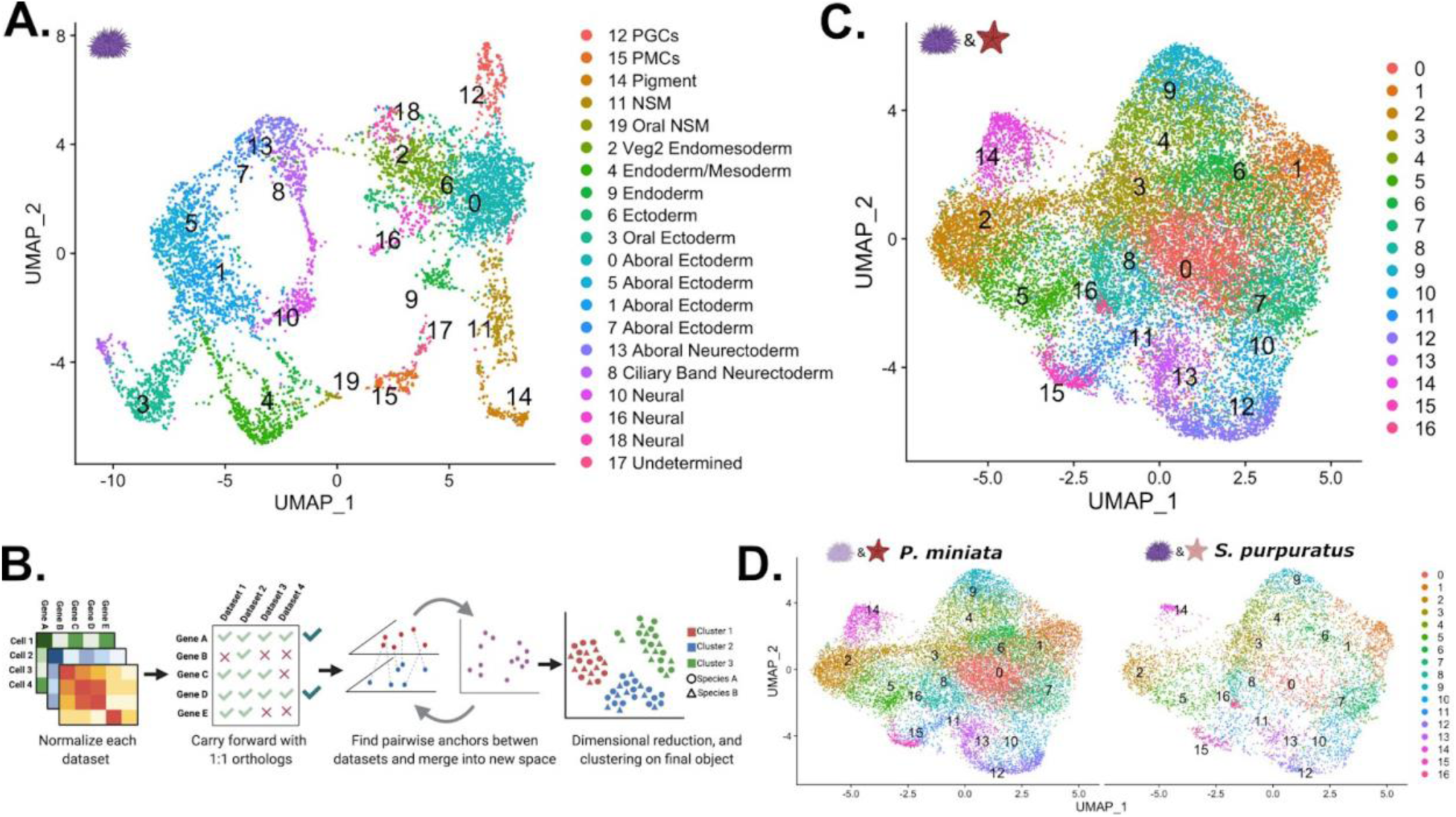
Creation of a developmental atlas in S. purpuratus and integration into a multi-species atlas. (A) UMAP projection of the 20 clusters identified and annotated in our *S. purpuratus* dataset used in later analysis. (B) Our pipeline for creating a multi-species atlas that uses only 1:1 orthologs to integrate the datasets into one common space using CCA. (C) The UMAP reduction of our multi-species atlas with 17 identified clusters. (D) Views of only the *P. miniata* (left) and *S. purpuratus* (right) data points in the multi-species atlas.

Pigment cells (Spur CL 14) and PMCs (Spur CL 15) were identified as distinct annotated clusters on the *S. purpuratus* UMAP projection (Figure 8.A). The PMC cluster was marked by the expression of both regulatory genes like *ets1* (adjusted p-value = 1.38E-77), aristaless-like homeobox (*alx1*, adjusted p-value =4.02E-300), T-box brain transcription factor 1 (*tbr*, adjusted p-value = 1.31E-118), ets variant transcription factor 6 (*etv6*, adjusted p-value = 1.12E-53), transcriptional regulator ERG (*erg*, adjusted p-value = 8.02E-231), hematopoietically-expressed homeobox protein HHEX homolog (*hhex*, adjusted p-value =1.37E-160), TGFB induced factor homeobox 2-like (*tgif2*, adjusted p-value = 6.96E-87), *arid3a* (adjusted p-value = 1.05E-138), forkhead box O1 (*foxo1*, adjusted p-value = 1.06E -15), and vascular endothelial growth factor receptor 1 (*flt1*, also known as *VEGFR1*, adjusted p-value = 0.00E+00) and genes involved in biomineralization, including 27 kDa primary mesenchyme-specific spicule protein (*pm27*, adjusted p-value = 5.66E-114), spicule matrix protein SM50 (*sm50*, adjusted p-value = 1.76E- 117), and mesenchyme-specific cell surface glycoprotein (*msp130*, adjusted p-value = 0.00E+00).

The pigment cell cluster had high expression of genes known to be involved in pigment cell specification. These include probable polyketide synthase 1 (*pks1*, adjusted p-value=9.90E-88), *gcml* (adjusted p-value=1.77E-44), growth factor independent 1 transcriptional repressor (*gfi1*, adjusted p-value=4.78E-65), *gata6* (adjusted p-value=8.58E-66), *gata3* (adjusted p-value=3.01E-35), prospero homeobox 1 (*prox1*, adjusted p-value=5.12E-53), and ets homologous factor (*ehfl*, adjusted p-value=6.02E-18).

### Multispecies integration

In order to directly compare sea star and sea urchin cell types, we projected the datasets from the two species into a single shared space (Figure 8.B). We took advantage of extensive orthology analysis that has identified 1:1 orthologs between *P. miniata* and *S. purpuratus*, using a DIOPT-like system (Foley et al., 2021) and subsetted the datasets to only include 1:1 orthologs that are expressed in both datasets (see methods). This filtered gene set had 5,790 genes (28.3% of *P. miniata’s* total genes, 21.7% of *S. purpuratus’* total genes), removing species-specific genes and genes with 1:many orthologs from our analysis. Normalization, integration, clustering, and dimensional reduction were done according to the same pipeline used in creating our single-species atlases and is detailed in the methods section. Our analysis resulted in 17 integrated clusters, (henceforth referred to with the prefix Int CL ID) (Figure 8.C-D).

### Integrated clusters show similar patterns of gene expression between species

We first questioned whether any of the integrated clusters were over-represented by cells/nuclei from one species. After normalizing for the number of cells/nuclei per species sample, we calculated the percentage of species-specific cluster cells/nuclei in each integrated cluster (Figure 9.A). For Int CL 0 through Int CL 15, each species contributes to at least a third of the total integrated cluster contents, except for Int CL 16, which is 92.9% *S. purpuratus* cells (A table listing all cluster contributors is available in Supplementary table 3). Therefore, we concluded that other than possibly Int CL 16, integrated clustering was not driven solely by species differences, batch effects, or any technical variations in the sampling method.

**Figure 9:**
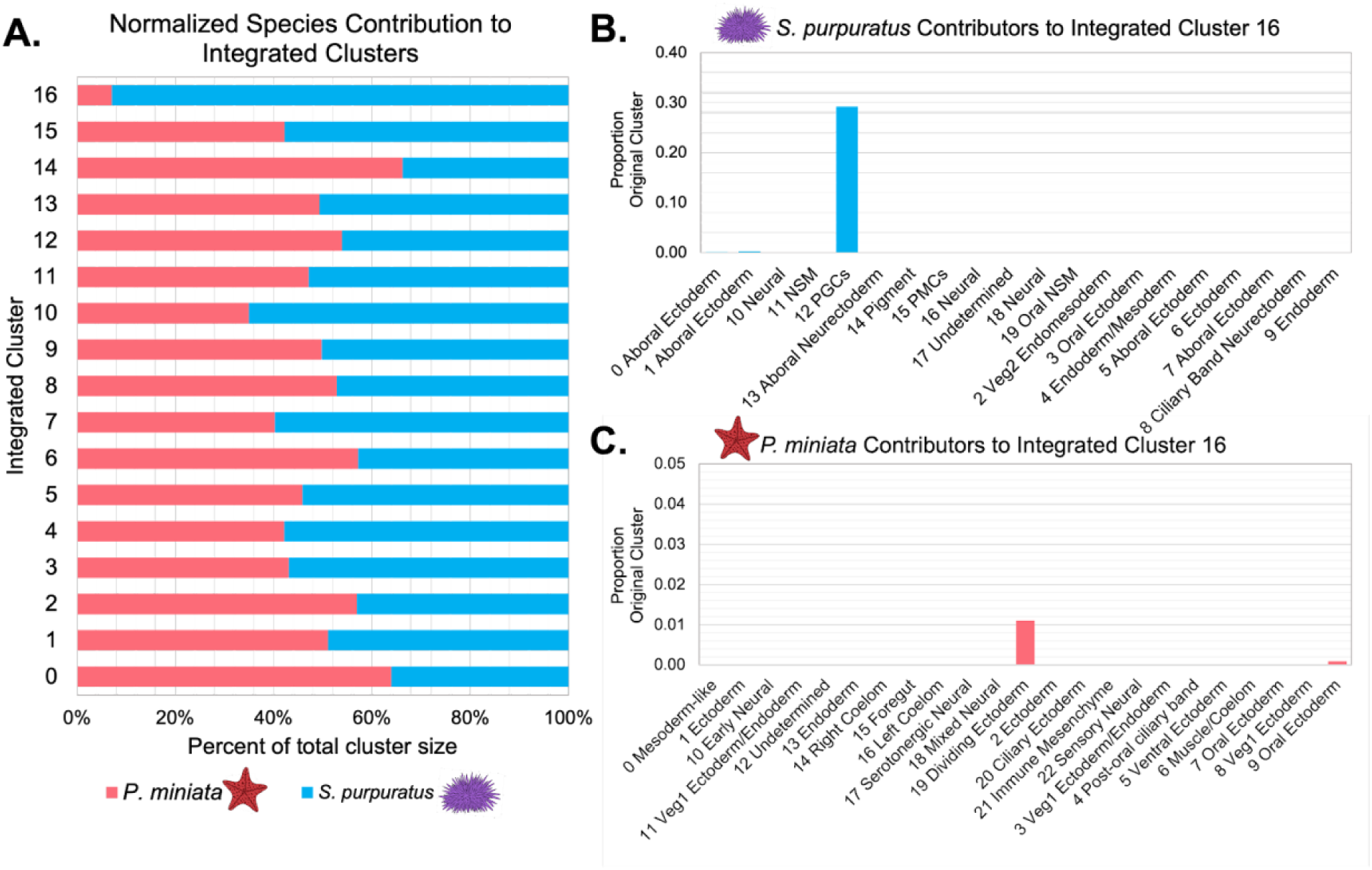
Quantification of species’ contribution to integrated clusters and the identification of a cluster enriched for *S. purpuratus* cells. (A) Bar graph shows the percentage of nuclei/cells of each species in each integrated cluster, normalized to account for differences in sample sizes. Species contribution is balanced in all but Int CL 16. (B) A bar graph showing the percentage of each Spur CL # that contribute to Int CL 16. (C) A bar graph showing the percentage of each Pmin CL # that contributes to Int CL 16.

To explore the compositions of these integrated groupings, we examined the contribution of the annotated clusters from our species-specific-atlas. For each integrated cluster, we calculated the percentage of each original cell type (Spur CL ID, Pmin CL ID) present in the integrated cluster (Int CL ID). A full table of these values is available in supplementary table 3.

### Primordial Germ Cells are an *S. purpuratus-*specific Integrated cluster

Int CL16 was the most species-specific integrated cluster, composed of 7.1% *P. miniata* nuclei (13 nuclei total) and 92.9% *S. purpuratus* cells (73 cells in total) (Figure 9.A). When we examine the composition of *S. purpuratus* cells present in Int CL 16 (Figure 9.B), we find that 70 of the cells originate from the Spur CL 12 Primordial Germ Cells (PGCs) (29.2% of all cells from Spur CL 12 PGCs), 2 cells originate from Spur CL 1 Aboral Ectoderm (2.44% of all cells from Spur CL 1 Aboral Ectoderm), and 1 cell from Spur CL 0 Aboral Ectoderm (0.08% of all cells from Spur CL 0 Aboral Ectoderm). The *P. miniata* contribution to Int CL 16 is made up of 12 nuclei from Pmin CL 19 Dividing Ectoderm (1.10% of all nuclei from Pmin CL 19 Dividing Ectoderm) and 1 nucleus from Pmin CL 9 Oral Ectoderm (0.09% of all nuclei from Pmin CL 9 Oral Ectoderm) (Figure 9.C).

We hypothesize that the very low membership of *P. miniata* nuclei in this integrated cluster indicates that there are no populations in *P. miniata* that closely resemble *S. purpuratus*’s PGCs and their presence in this cluster is likely driven by statistical noise and not transcriptomic similarity, thus making Int CL 16 the only *S. purpuratus*-specific cell type at this stage of development.

Of the other cells in Spur CL 12 PGCs, most (43% of all cells from Spur CL 12 PGCs) segregate into Int CL 8. Int CL 8 also contains 38.4% of all of *P. miniata*’s Pmin CL 19 Dividing Ectoderm, however, this only makes up 11.6% of the *P. miniata* nuclei in Int CL 8. The majority of the nuclei in this integrated cluster originate from 20 different Pmin CLs, with the maximum contribution being 13.29% of Pmin CL 12 Undetermined. Because this cluster is made up of such an amalgamation of *P. miniata* nuclei, we conclude their clustering is, at least in part, driven by similarities in the expression of cell cycle genes and not transcriptional regulatory genes.

Differences in the timing of PGC formation is consistent with previous work, which has shown that the genes typically used to identify PGC identity, such as *Piwi*, *Nanos*, and *Pumillo*, only begin to show cell-type-specific expression in *P. miniata* by the late gastrula stage when the posterior endocoelom forms (Fresques et al., 2014), a time point that lies outside our sampling window. Therefore, this cluster can be considered novel to *S. purpuratus* in our sampled window. The identification of this cluster demonstrates our method of atlas comparison allows for the detection of species-specific clusters.

### *S. purpuratus* pigment cells share transcriptomic features with *P. miniata*’s immune mesenchyme and neurons

Despite the novelty of their phenotype, and their distinct clustering pattern in the *S. purpuratus* cell atlas, the pigment cell cluster (Spur CL 14) does not form a novel transcriptomic cluster in the multispecies integrated clustering. Cells originating from Spur CL 14 pigment cells were primarily distributed between Int CL15 (40.1%) and Int CL 12 (26.7%) (Figure 10.A). Pigment cells also contributed to Int CL 2 (8.02%), Int CL 11 (7.49%), Int CL 5 (7.49%), Int CL 13 (6.95%), Int CL 10 (2.14%), Int CL 6 (0.54%), and Int CL 1 (0.54%) (Figure 10.A).

**Figure 10:**
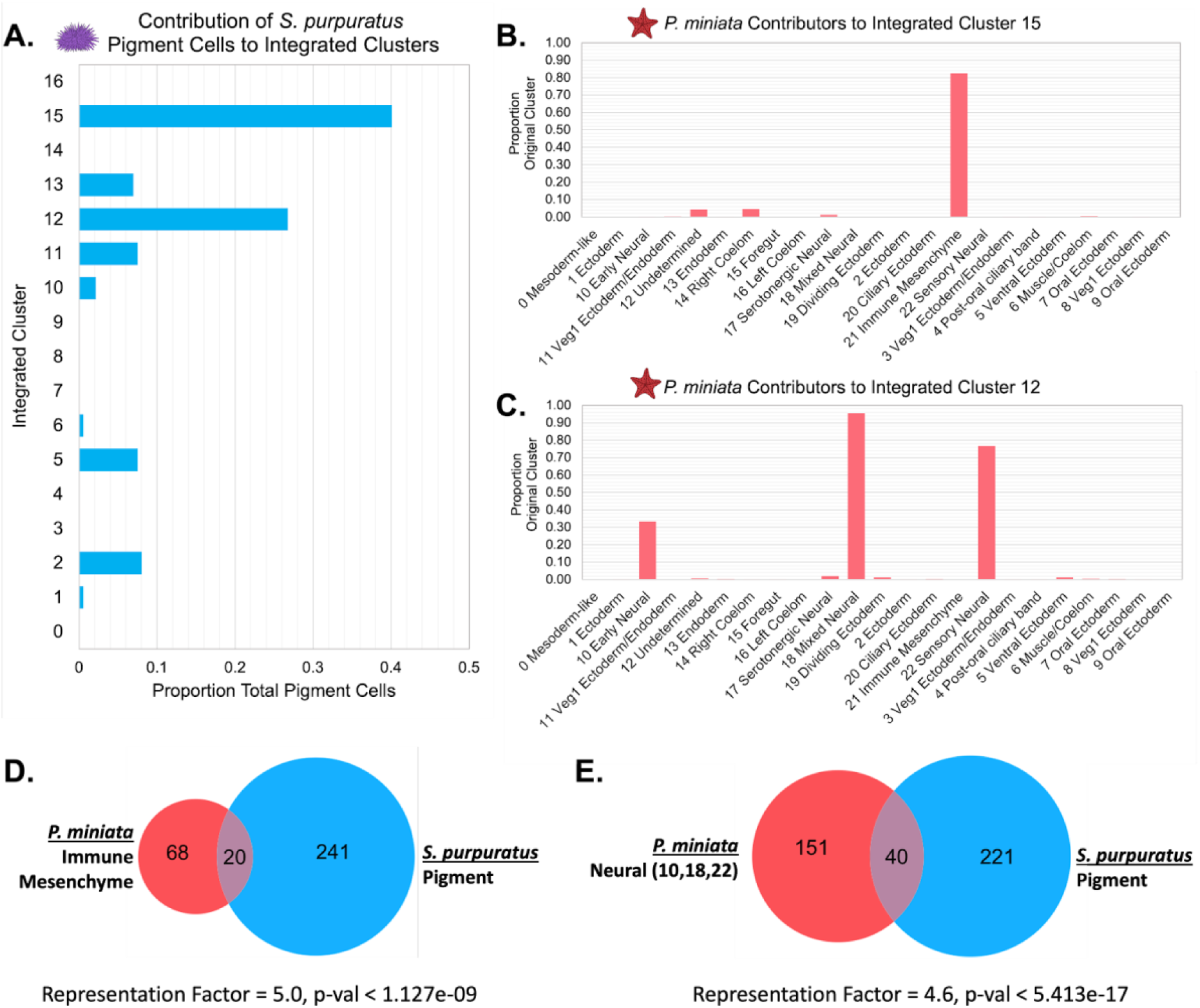
Pigment cells share transcriptomic similarities with *P. miniata* immune cells and neurons. (A) The proportion of total pigment cells from the *S. purpuratus* atlas that contribute to each Int CL. The pigment cells chiefly contribute to Int CL 15 and Int CL 12. (B) A bar graph showing the percentage of each Pmin CL # that contribute to Int CL 15. *P. miniata* cells in Int Cl primarily originate from Pmin CL 21 immune mesenchyme. (C) A bar graph showing the percentage of each Pmin CLs that contribute to Int CL 12. The *P. miniata* cells in Int Cl 12 primarily originate from three neural clusters: Pmin CL 10 early neural, Pmin CL 18 mixed neural, and Pmin CL 22 sensory neural. . (D) A Venn diagram highlights marker genes that are shared between the Spur CL pigment and Pmin CL immune mesenchyme. The overlap is higher than expected by random, with a representation factor of 5.0 and p-value < 1.127e-09. (E) A Venn diagram highlights the marker genes shared between Sur CL pigment and the pooled markers genes of Pmin CL 10 early neural, Pmin CL 18 mixed neural, and Pmin CL 22 sensory neural. The representation factor of this overlap is 4.6 and p-value < 5413e-17.

Pmin CL 21 immune mesenchyme contributed 82.5% of nuclei of its original annotation to the Int Cl 15 (Figure 10.B). There were also smaller contributions from Pmin CL 14 Right Coelom (4.60%), Pmin CL 12 Undetermined (4.26%), Pmin CL 17 Serotonergic Neurons (1.10%), Pmin CL 5 Endoderm/Coelom (0.52%), Pmin CL 11 Veg1 Ectoderm/Endoderm (0.3%), Pmin CL 13 Endoderm (0.19%), Pmin 10 Neural (0.12%), Pmin 8 Veg1 Ectoderm (0.11%), Pmin CL 3 Veg1 Ectoderm/Endoderm(0.09%),and Pmin CL 4 Post-oral Ciliary Band (0.09%).

When comparing the marker genes of the individual species atlas, we see 20 1:1 orthologous marker genes shared between the Pmin CL immune mesenchyme and Spur CL pigment cell (Figure 10.D). To examine the statistical significance of the overlap in marker gene sets, we calculated the representation factor based on the overlap of 1:1 orthologs between clusters across all 1:1 orthologs. *P. miniata* immune mesenchyme and *S. purpuratus* pigment cells have a statistically significant overlap in markers, with a representation factor of 5.0 and corresponding p-value < 1.127e-09. Amongst these genes were *Pax6*, which has been previously described to play a role in sea urchin mesoderm formation (Martik and McClay, 2015) and *gata3*, which has been shown to play a role in coelomocyte formation in *S. purpuratus (Pancer et al., 1999)*. They also share genes with characterizing immune-responsive functions in other animals, including *Srcr* and *rap2c* (Gillespie et al., 2022; Liu et al., 2011). We, therefore, predict that *S. purpuratus* pigment cells and *P. miniata* immune mesenchyme share conserved immune response elements.

Secondly, *P. miniata*’s contributors to Int CL 12 primary arise from three neural populations - Pmin CL 10 Neurons (33.4%), Pmin CL 18 Mixed Neural (95.6%), and Pmin CL 22 Sensory Neural (76.7%). We also see small contributions from Pmin CL 17 Serotonergic Neurons (1.93%), Pmin CL 5 Endoderm/Coelom (1.20%), Pmin CL 19 Undifferentiated Ectoderm (1.13%), Pmin CL 12 Undetermined (0.68%), Pmin CL 6 Mesoderm/Coelom (0.62%), Pmin CL 20 Ciliary Ectoderm (0.39%), Pmin CL 14 Endoderm (0.38%), Pmin CL 7 Oral Ectoderm (0.33%), Pmin CL 1 Ectoderm (0.27%), Pmin CL 15 Foregut (0.20%), Pmin CL 14 Right Coelom (0.19%), Pmin CL Veg1 Ectoderm/Endoderm (0.18%), Pmin CL 4 Post-oral Ciliary Band (0.18%), Pmin CL 2 Ectoderm (0.14%), and Pmin CL 8 Veg1 Ectoderm (0.11%) (Figure 10.C).

Comparisons of the marker gene sets of the pigment cells and merged neural cells (excluding serotonergic cells), identified 40 gene orthologs in common, with a representation factor of 4.6 and corresponding p-value < 5.413e-17 (Figure 10.E). These clusters had several transcription factors shared as marker genes: *foxn3*, homeobox protein Hox-D9a (*Hox*-*D9a*), ETS variant transcription factor 1 (*etv1*), and *gcml*. In *S. purpuratus*, gcml is a known marker of pigment cells (Calestani et al., 2003),(Ransick and Davidson, 2012). Several other genes related to neuronal function were also identified as shared markers including *SYT14*, *thrb*, 3 voltage-gated channels (voltage-dependent calcium channel subunit alpha-2/delta-1 (*CACNA2D1*), non L-type Cav channel(*CcLta1*), and voltage-dependent L-type calcium channel subunit beta-1 (*CACNB1*)), and rab3 GTPase (*rab3*). *P. miniata* neurons and *S. purpuratus* pigment cells also share marker genes potentially involved in immune function, including *Srcr* and nemo-like kinase (*nlk*) (Lv et al., 2016)

### *S. purpuratus*’ PMCs cluster with *P. miniata*’s foregut and right coelom

Spur CL 14, annotated as PMCs, do not form a distinct transcriptomic cluster in the multi-species integration map (Figure 11.A). However, 60.3% of all PMCs segregate into Int CL 5, while the rest are scattered across different integrated clusters. Thus PMCs also contribute low percentages of cells to Int CL 13 (10.3%), Int CL 6 (5.13%), Int CL 0 (4.49%), Int CL 3 (4.49%), Int CL 14 (4.49%), Int CL 2 (3.21%), Int CL 11 (2.56%), Int CL 10 (1.92%), Int CL 7 (1.28%), Int CL 15 (1.28%), and Int CL 8 (0.641%).

**Figure 11:**
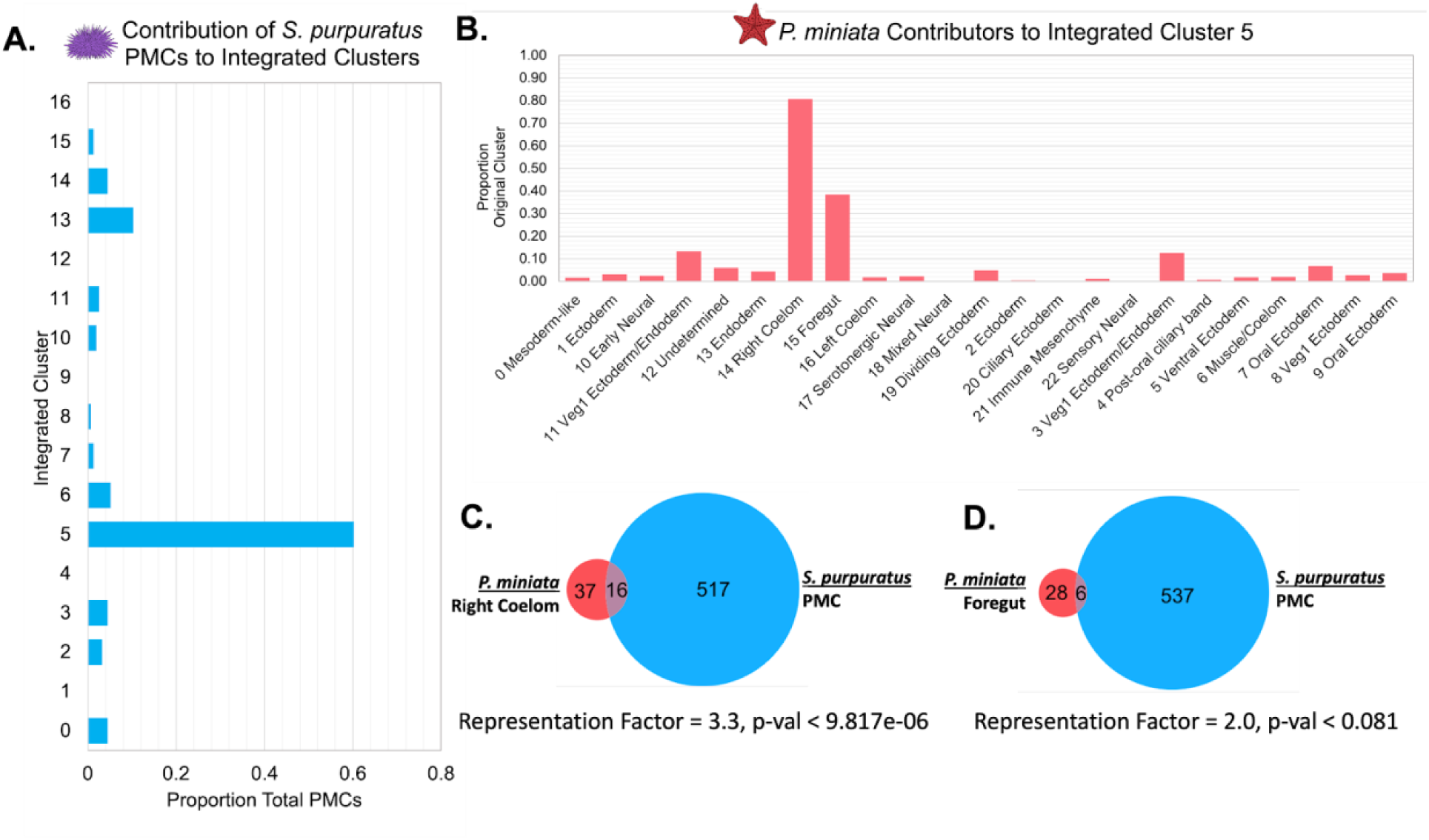
PMCs share transcriptomic features with *P. miniata*’s right coelom and foregut. (A) The proportion of total PMC cells from the *S. purpuratus* atlas that contribute to each Int CL. More than half of all PMCs segregate into Int CL 5. (B) A bar graph showing the percentage of each Pmin CL # that contribute to Int Cl 5. The top contributors are nuclei from Pmin CL 14 Right Coelom and 15 Foregut. (C) Venn diagram highlights the overlap of marker genes that are 1:1 orthologs that are shared between *S. purpuratus*’ PMCs and *P. miniata’*s Right Coelom. The representation factor is 3.3, with p-value < 9.817e-06. (D) Venn diagram highlights the overlap of marker genes that are 1:1 orthologs between *S. purpuratu*s’ PMCs and *P. miniata*’s foregut. The representation factor is 2.0 with p-value < 0.081.

When we examine *P. miniata* cell types contributing to Int CL 5, the top contributors are Pmin CL 14 Right Coelom (80.7%) and Pmin CL 15 Foregut (38.4%) (Figure 11.B). There were also low-level contributions from most other *P. miniata* clusters, with representatives from Pmin CL 11 Veg1 Ectoderm/Endoderm (13.3%), Pmin CL 3 Veg1 Ectoderm/endoderm (12.6%), Pmin CL 7 Oral Ectoderm (6.84%), Pmin CL 12 Undetermined (5.96%), Pmin CL 19 Undifferentiated Ectoderm (4.91%), Pmin CL 13 Endoderm (4.37%), Pmin CL 9 oral ectoderm/foregut (3.78%), Pmin CL 1 Ectoderm (3.12%), Pmin CL 8 Veg 1 Ectoderm (2.71%), Pmin CL 10 Neurons (2.51%), Pmin CL 17 Serotonergic Neurons (2.20%), Pmin CL 6 Mesoderm/Coelom (1.98%), Pmin CL 16 Left Coelom (1.79%), Pmin CL 5 Endoderm/Coelom (1.76%),Pmin CL 0 Mesoderm (1.67%), Pmin CL 21 Immune Mesenchyme (1.19%), Pmin CL 4 Post-oral Ciliary Band (0.73%), and Pmin CL 2 Ectoderm (0.49%).

A comparison of *P. miniata’s* foregut markers to *S. purpuratus’* PMC markers yielded a representation factor of 2.0 with a p-value < 0.081 while the overlap of markers between *P. miniata’s* right coelom and the PMCs resulted in a representation factor of 3.3 and p-value < 9.17e-06 (Figure 11.C-D). The 4 genes shared across all 3 original clusters are *fibrillin-3,* sprouty-related EVH1 domain-containing protein 3 (*spred2*), 3 alpha procollagen precursor (*col4a1*), and eukaryotic translation initiation factor 4 gamma 3 (*EIF4G3*). Both serine/arginine repetitive matrix protein 1 (*SRRM*) and probable JmjC domain-containing histone demethylation protein 2C (*HNHD1C*) act as markers in only *S. purpuratus*’ PMCs and *P. miniata* foregut.

In total, 12 marker genes are shared between the PMCs and *P. miniata*’s right coelom. Of particular interest are the shared markers, *arid3a*, transcription factors *Elk*, *foxn3,* and *ets1*, genes previously characterized in the PMC GRN of *S. purpuratus* (Davidson et al., 2002b; Rho and McClay, 2011). This suggests there is a region of *P. miniata* that directly corresponds to PMCs, despite morphologically lacking them.

### *P. miniata* left coelom has no equivalent *S. purpuratus* cell cluster

Unlike the right coelom, there is little overlap in marker gene content between the left coelom and PMCs. These two groups share 4 genes in common (solute carrier family 4 member 11 (*slc4a110*), 3 alpha procollagen precursor (*col4a1*), eukaryotic translation initiation factor 4 gamma 3 (*EIF4G3*), and Rap associating with DIL domain (*radil*), with a representation factor of 1.6 and p-value < 0.229, making it statistically insignificant.

We, therefore, were especially interested in comparing the cluster identity of the left coelom. 95% of Pmin CL 16 Left Coelom nuclei contribute to Int CL 14 (Figure 12.A). When we examine the contributions of *S. purpuratus* cells to that cluster, we see that there is no main contributor, rather small contributions across a wide range of *S. purpuratus* cell types, originating from 16 different Spur CL clusters, with contributions ranging from 0.54% to 4.49% of a Spur CL. Spur CL 15 PMCs (4.49%), Spur CL 13 Aboral Neurectoderm (3.14%), Spur CL 8 Ciliary Band Neurectoderm (2.97%) Spur CL 5 Aboral Ectoderm (2.56%), Spur CL 2 Veg2 Endomesoderm (2.54%), Spur CL 16 Neural (2.54%), Spur CL 19 Oral NSM (2.34%), Spur CL 3 Oral Ectoderm (2.23%), Spur CL 11 NSM (2.21%), Spur CL 10 Neural (2.16%), Spur CL 9 Endoderm (2.11%), Spur CL 7 Aboral Ectoderm (1.77%), Spur CL 0 Aboral Ectoderm (1.76%), Spur CL 4 Endomesoderm (1.52%), and Spur CL 6 Ectoderm (0.542%) make up the *S. purpuratus* portion of Int CL 14.

**Figure 12:**
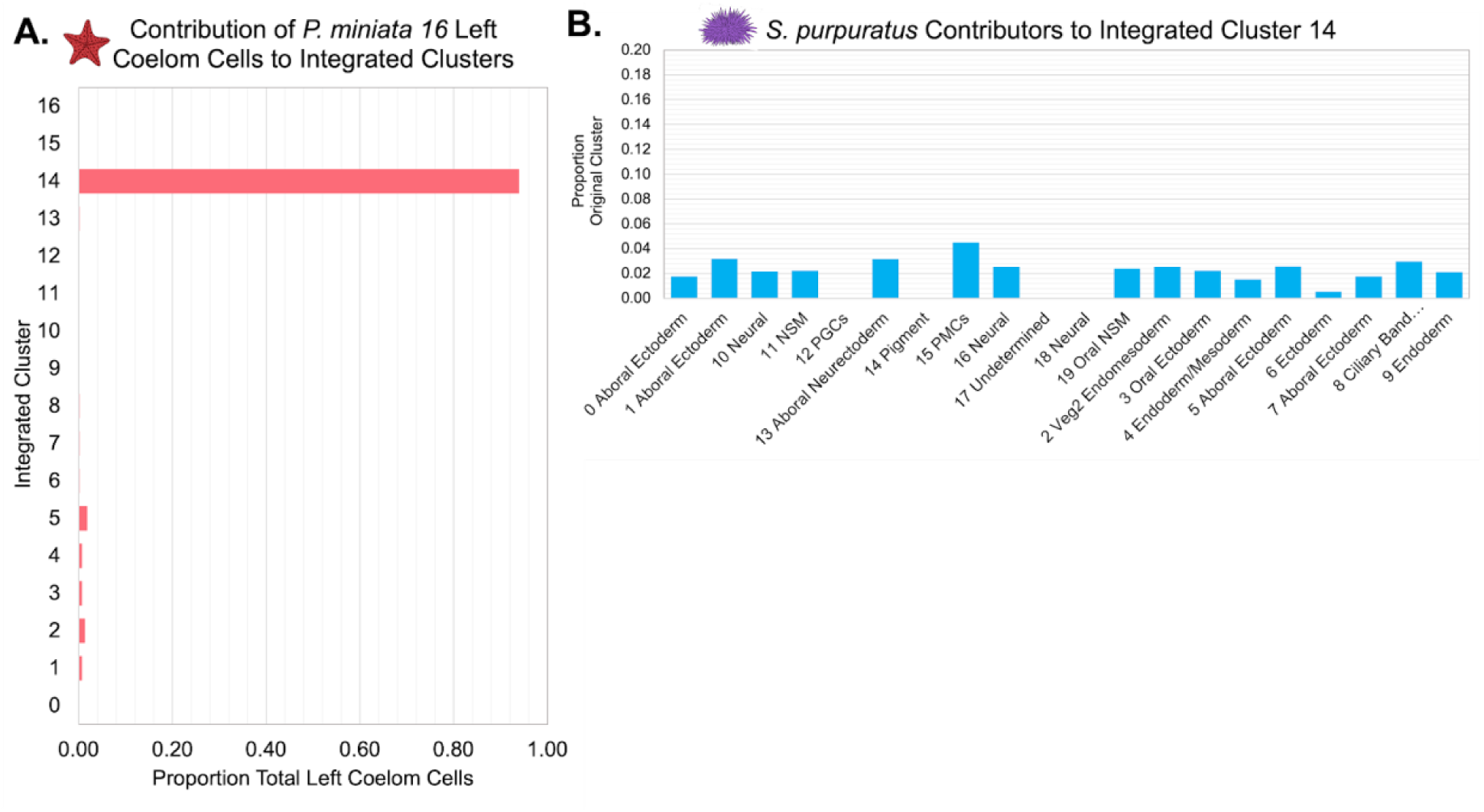
The *P. miniata* left coelom does not have a consistent corresponding population in *S. purpuratus.* (A) The proportion of total nuclei originating from Pmin CL 16 Left Coelom across all integrated clusters. Most of the nuclei are located in Int CL 14. (B) A bar graph showing the percentage of each Spur CL # that contributed to Int CL 14. The cells in Int CL 14 originate from many different Spurp CL # and no single cluster contributes more than 4.49% of a cluster’s total cells.

There are several potential explanations for the isolation of the left coelom cells in our integrated clustering. Firstly, this could correspond to a novel cell lineage that has not been documented in sea stars. Alternatively, the transcriptomes of homologous coelomic pouches could be sufficiently divergent so that they no longer drive clustering in the integration approaches we used, possibly relating to the morphological uniqueness of sea star coeloms with respect to other echinoderm species (Child, 1941; Morris et al., 2009). Secondly, this could, like with the PGCs, result from a difference in developmental timing, and the corresponding population has not yet arisen in *S. purpuratus* at the sampled stages. Finally, this clustering could have arisen as an artifact of sampling and/or clustering, not reflective of biological meaning. Further interrogation of *P. miniata’s* left coelomic pouch development is needed to better understand these phenomena.

## Discussion

At the heart of evolutionary developmental biology rest two questions: what is a cell type? And what is novelty? Being able to differentiate and reproducibly define cell categories is essential to understanding the evolutionary relationship between cell types and the cross-species comparisons needed to establish novelty. The identification and definition of cell types can change with new technologies as new data is brought to light. In this paper, a single-nucleus RNA-sequencing approach was used to classify cell types present during early embryonic development in the bat star, *P. miniata*. An integrated atlas was constructed using 1:1 orthologs of *P. miniata* and *S. purpuratus* to investigate and infer the origins of novel cell types in these species.

### An early embryo developmental atlas for *P. miniata*

We produced the first developmental atlas of an echinoderm belonging to the Asteroidea family. The expected cell types were observed and known markers of major tissue types were identified (summarized in Figures 3 and 4). We were also able to make several key observations of cell types present in sea star embryos for the first time. We observed 4 neural clusters in our analysis. These include serotonergic neurons, sensory neural, mixed neural, and an early, general neural type. The identification of these distinct neural clusters is consistent with the finding that sea urchin early gastrula embryo has three neural cell types (Slota and McClay, 2018). In sea urchins, these three neural subtypes are distinguished by the expression of unique transcription factors. The work here shows that neural subtypes can also be distinguished by the expression of different synaptotagmin genes in *P. miniata*, suggesting distinct synaptic vesicle trafficking in these neural types. The identification of endocrine-associated markers in two of these neural clusters (Pmin CL 18 Mixed Neural and Pmin CL 22 Sensory Neural) supports an emerging hypothesis of the origins of neural subtypes from secretory cells (Moroz, 2021), which is thus still evident in the mixed endocrine and neural markers of these clusters. This is also consistent with recent work in *S. purpuratus* larvae, which identified endocrine-associated genes in post-oral/lateral neurons and argues that the endocrine system evolved from a subset of neurons (Paganos et al., 2021; Perillo et al., 2018). The similarities found in sea stars and sea urchin embryos demonstrate that the striking degree of neural diversity is a likely feature of early eleutherozoan embryos.

Diverse mesodermal cell types were also identified by snRNA-seq in *P. miniata*, including an immune population (Figures 4 and 6). In the *P. miniata* immune mesenchyme cluster were orthologs of the sea urchin blastocoelar and pigment cells’ immune response gene *tek*, a gene expressed in sea urchin coelomocytes and linked to the proliferation of immune cells following an immune challenge (Stevens et al., 2010). *P. minata* immune mesenchyme and sea urchin blastocoelar cells also share the marker genes *duox1* and mitogen-activated protein kinase 1 (*mapk1*), which are well documented to play a role in stress and immune response (Zhan et al., 2018); (Pinsino et al., 2015; Smith et al., 2018). These genes are not detected as markers in the sea urchin pigment cell cluster, suggesting that the pigment and blastocoelar cells play different roles in the immune response.

The two populations of coelomic mesoderm in *P. miniata* are of special interest. The right coelom shares many markers with the immune mesenchyme cluster (including transcription factors such as *elk* and *ets1*, Figures 4 and 6), suggesting that the immune mesenchymal cells originate from the right coelom and may continue to do so over development. However, the left coelom does not have mesenchyme markers suggesting that cells in the left coelom do not undergo epithelial to mesenchymal transition at this developmental stage. This asymmetry is maintained throughout later developmental stages outside of our sampling window (e.g. shown by whole-mount in situ hybridization in Figure 7.D.iv-v). This for the first time, suggests a very distinct role of the left and right coeloms in sea stars.

### A comparative single nuclei/cell transcriptomic approach for studying the evolution of cell types

Identifying homologous cell types across species is a central quest in biology as it is the prerequisite knowledge for understanding the evolutionary basis of cell and morphological diversification.

Here, we have presented an orthology-based methodology for combining developmental single cell/nuclei data across species into one composite atlas. Current approaches focus on either integrating data from multiple species into a common atlas or performing pairwise comparisons between separately clustered datasets (Shafer, 2019; Tanay and Sebé-Pedrós, 2021). An advantage of examining clustered multi-species transcriptome data is that it accommodates split clustering of an organism’s original cluster identity, where a transcriptomic profile defined in a species-specific developmental atlas may split between more than one cluster into the integrated data. This allows us to pull out elements of the GRN that overlap, which we can combine with biological knowledge to form hypotheses for the sister cell relationships and the identification of shared modules deployed in different contexts. We have shown that using only 1:1 orthologs to create an integrated atlas is sufficient to drive clustering that results in mixed-species clusters that demonstrate meaningful transcriptomic similarity. However, it should be noted that restricting our analysis to only 1:1 orthologs removes data from an analysis that can potentially mask the importance of non-orthologous genes and gene duplications in driving species-specific cell-type differences. As this field advances, integration accounting for paralogous transcripts could help improve the meaningfulness and depth of transcriptome comparisons.

To explore the concept of cell novelty, we first examined whether any integrated clusters showed a species bias. If a cell type was completely unique to one species, we would expect to see a cluster on our integrated UMAP that is dominated by contributions from one species. Integrated cluster 16 is our most species skewed cluster, with 7.1% from our *P. miniata* dataset and 92.9% of its members originating from our *S. purpuratus* dataset, most of which were originally annotated as PGCs (Figure 10). The sea urchin PGCs could therefore be viewed as novel under the GRN definition of cell types as they do not share any significant transcriptional similarity with cell clusters in sea stars. However, an alternative interpretation is that these cells have yet to be specified in the stages of our sampling in sea stars. There is evidence that these cells are specified later in sea stars (Fresques et al., 2014). Therefore, we propose that PGCs can be considered either novel or temporally novel to *S. purpuratus* in our dataset, driven by differences in developmental timing and not separate evolutionary cell lineages. This distinction can be resolved by expanding the datasets to later time points and establishing whether the PGCs would form mixed-species clusters.

Our analysis also identified another potentially species-specific cell population, the left coelom of *P. miniata*, which primarily contributed to Int CL14 (Figure 10). Though it was not species-exclusive, the *S. purpuratus* atlas of the cells that contributed to the integrated cluster were of numerous identities with no clear contribution from any cluster in *S. purpuratus*. In both species, the left coelomic pouch gives rise to the hydrocoel, which forms the adult water vascular system in adults and is a shared feature of all echinoderms (Balser et al., 1993; MacBride, 1918). Therefore from a functional and developmental perspective, these cells are not considered novel, but using our definition of transcriptome similarity they may represent a novel population in sea stars. Understanding the GRN operating in this coelom will provide further insights into the role of this unique regulatory transcriptome.

In comparison, the presumed, traditionally defined novel cell types, i.e. the PMCs and pigment cells in sea urchins form molecularly distinct populations in the single species atlas, but lose that distinctness in our multi-species atlas where we find similarities between their transcriptomes and those of *P. miniata* clusters. In our integrated data set, the PMCs aggregate with nuclei from *P. miniata*’s right coelom and these two populations share a significant number of marker genes (Figure 12), including several transcription factors (ets1, dri, elk, foxn3, and spred2). We suggest that these genes have a conserved role in the core mesenchyme specification program present in their last common ancestor.

Of particular interest is our detection of *VEGFR1L* in *P. miniata*. Previous studies have shown that ortholog of this gene, *VEGFR1* (also known as *flt1)* was exclusively expressed in the skeletogenic mesenchyme of brittle stars and sea urchins and was responsible for the deployment of the biomineralization gene set as was seen as a critical node of cooption (Morino et al., 2012). *VEGFR* signaling does not activate skeletogenesis pathways, at least at this developmental stage, as the sea star embryo does not make a skeleton. The *P. miniata* PMC- like GRN now identified in the right coelom may instead have a conserved role in cell guidance, and/or establishing the clustered ring of primary mesenchymal cells at the bottom of the blastocoel, as it does in skeletogenic-PMCs (Morino et al., 2012). In Figure 6.G., we see *VEGFR* positive cells form a ring around the base of the archenteron, similar to skeletogenic PMC cells. *VEGFR* has also been shown to control conserved tubulogenesis programs documented in other echinoderms and vertebrates (Ben-Tabou de-Leon, 2022; Morgulis et al., 2019). The coelomic pouches of asteroids are very distinct from other species of echinoderms; they form much earlier in embryonic development and are longer and thinner, compared to other echinoderms (Child, 1941). Therefore, we hypothesize that some of those conserved tubulogenesis genes may also be involved in developing their distinct coelomic morphologies.

In our multi-species atlas, pigment cells that formed a cluster in the *S. purpuratus* atlas, contribute to integrated clusters with *P. miniata* immune mesenchyme or gcml+ neurons. It is not surprising that pigment cells may share similarities with immune cell types in sea stars, given their presumed shared immune functions. When comparing the immune mesenchyme of the sea star with the pigment cells of *S. purpuratus* we see several shared combinations of transcription factors, previously demonstrated to play similar roles in cell fate specification between *S. purpuratus* and *P. miniata*. The sharing of these early-acting regulators suggests that there is a common mesodermal and immune mesenchyme regulatory program maintained between *S. purpuratus* and *P. miniata*.

The pigment cell clustering with *P. miniata* neurons, however, was unexpected. When comparing the gene expression patterns of the pigment cells and *P. miniata* neurons, we see an overlap of several genes. Of particular interest is their shared expression of *gcml*. Previous literature showing *gcml* is expressed in neural development in other phyla (Jones et al., 1995; Soustelle et al., 2007) leads us to suggest that gcml plays a role in the development of neural-like excitable or secretory cells in echinoderms, and in the common ancestor of the eleutherozoa. In this scenario, pigment cells may have arisen through the co-option of a neuronal *gcml* module in a subset of mesenchyme cells. The regulatory independence of this new *gcml*-expressing population allowed for genetic individualization and the evolution of different downstream functionalities including the activation of pigment formation genes. At the same time, we see the maintained expression of other neuronal effector genes (*SYT14*, *thrb*, voltage-gated channels, and *rab3*), possibly another consequence of *gcml* expression. This suggests that these genes may play dual roles in immune and neural cells.

## Conclusion

We have used single nuclei RNA-seq to study early embryo development in the sea star *P. miniata* and an orthology-centered approach to integrate it with the atlas of sea urchin, *S. purpuratus.* Single-cell transcriptomics provides a very detailed picture of gene expression and cell identity. This allows us to deeply explore concepts of cell-type identity and evolutionary novelty through the lens of gene expression and inferred GRNs operating in the cells. Our comparisons discovered examples of transcriptomically distinct cell clusters, i.e. PGCs in sea urchins and left coelom in sea stars, with no clear orthologous cluster in the other species. These may be taken as examples of a novel cell when defined on the basis of gene expression similarities achieved by this method. Novelty in both of these cases may be relative changes in timing or heterochrony of the cell state specification. Conversely, the two cell types that we anticipated as novel based on cell morphology, function, and developmental lineage, i.e. PMCs and pigment cells in sea urchins had significant transcriptomic profiles in common with sea star cell clusters. This implies a shared ancestry of these cells, with an ancestral cell possessing these regulatory profiles. In this case of PMCs and right coelom, this may include overlapping functions in EMT, cell migration, and tubulogenesis with a cooption, quite late in the developmental GRN of cassettes of biomineral genes. The most surprising find from our studies was the overlap between pigment cell cluster transcriptome and the immune and neural clusters in sea stars. This implies a more complex evolutionary scenario whereby programs from two cell types in sea stars have merged into one to produce a distinct cell morphology and function.

In conclusion, our study provides new interpretations of cell type novelty and new inferences of novel cell type evolution when considering comparisons of orthologous gene expression profiles over traditional approaches.

## Methods

### Embryo isolation and prep

Adult *P. miniata* and *S. purpuratus* animals were obtained by Marinus Scientific (Long Beach, California) and housed in 100-gallon tanks containing aerated artificial seawater (Instant Ocean, 32 – 33 ppt between 10-15°C).To induce spawning, adults were injected with 200 µM 1-methyl- adenine (Kanatani, 1969). Male and female gametes were harvested and mixed for 10 minutes to facilitate fertilization. Zygotes were filtered out using a 100µm mesh filter and washed with artificial seawater to remove excess sperm. Cultures were then transferred to 4-gallon buckets and incubated at 13-16°C, with constant gentle shaking to prevent settling.

### Whole-cell dissociation

The whole-cell dissociation protocol was adapted from (Barsi et al., 2014). Prior to dissociation, whole embryos were washed and resuspended in 500µL artificial seawater containing 2mg/mL Pronase, incubated on ice for 3 min, and centrifuged at 700g for 5 min at 4°C before being resuspended in ice-cold calcium-free artificial seawater (20mg/mL: BSA). The embryos were then manually dissociated using a P1000 micropipette and filtered through a 30 µM mesh filter. The resulting cells were centrifuged at 300g for 5 minutes at 4°C, and resuspended in artificial seawater.

### Nuclear isolation

Prior to dissociation, embryos were washed twice with cold seawater by centrifuging for 5 min (22g) at 4°C, and resuspended in 10 mL cold HB Buffer (15mM Tris pH7.4, 0.34 mM sucrose, 15mM NaCl, 60 mM KCl, 0.2mM EDTA, 0.2mM EGTA, 2% BSA, 0.2U/µL protease inhibitor (Sigma Aldrich cOmplete protease inhibitor)) and kept on ice. Washed embryos were then transferred to a 15mL dounce, and homogenized 20 times with the loose pestle and 10 times with the tight pestle. The dounced sample was centrifuged for 5 min (3500g) at 4°C, and the pellet was resuspended in 1mL HB buffer. Tip strainers (Scienceware® Flowmi™Flowmi) were used to transfer the sample to a new tube and 9mL ice cold HB buffer was added to the sample and centrifuged for 5 min (3500g) at 4°C. The pellet was resuspended in 1 mL PBT (1X PBS, 0.1% Triton X-100, 2% BSA, 0.2U/µL protease inhibitor (Sigma Aldrich cOmplete protease inhibitor tablet), and 0.2 U/µL SUPERase-in RNAse inhibitor (Invitrogen)). Samples were centrifuged for 5 min (2.4g) in a cold room and nuclei were resuspended in 300 µL PBT. Nuclei were stained with Trypan Blue and counted in triplicate with a Countess II Automatic Cell Counter (ThermoFisher). The resulting nuclei were resuspended to 1000 nuclei/µL.

### Bulk RNA-seq sample preparation and sequencing

Bulk RNA sequencing was performed on 25hpf *P. miniata* embryos that had been left whole, dissociated into single cells, or were nuclear isolates (Figure 2.A). Bulk RNA was isolated using the GenElute™ Mammalian Total RNA Miniprep kit (Sigma). Illumina HiSeq1 Libraries were prepared for Illumina HiSeq 51bp paired-end sequencing and sequenced on the NovaSeq 6000 (Duke Center for Genomics and Computational Biology, Durham, NC). Reads were aligned to the *P. miniata* genome version 2.0 (Kudtarkar and Cameron, 2017) using the standard TopHat pipeline (Kim et al., 2013), and gene counts were quantified using HTseq (Putri et al., 2021) and quantile normalized.

### Single nuclei library preparation and sequencing

Single nuclei libraries were created using the Chromium Next GEM Single Cell 3’ Reagent Kit v3.1 RevD (10X Genomics) and the Chromium X instrument (10X Genomics) using manufacturer protocols. For *S. purpuratus*, the number of nuclei recovered was 2,000 for the 6 hours post fertilization (hpf) sample, 5,000 for the 15 hpf and 6,000 for 23 hpf samples. For *P. miniata,* there were 3,000 nuclei in the 16hpf sample, 5,000 in the 26 hpf sample, and 6,000 in the 33hpf sample. After constructing the libraries, samples were quantitated on a Tapestation using D5000 ScreenTape® to confirm insert sizes of 400-500bp. Samples were sequenced on a Novaseq 2X150 S4 lane (Duke Center for Genomics and Computational Biology, Durham, NC).

### Processing and Quality Control of snRNA-seq data

Reads were processed using Cell Ranger V4.0.0 (Zheng et al., 2017) and aligned to a pre- mRNA index based on the *P. miniata* genome V3.0 (GenBank assembly accession: GCA_015706575.1) and *S. purpuratus* genome V5.0 (GenBank assembly accession: GCA_000002235.4). The pre-mRNA indices were constructed using gene annotation files modified by an in-house script to remove intron annotations, allowing unspliced reads to align. All further analysis was conducted in R V4.0.5 (R Core Team, 2021).

Quality control for our nuclear datasets was carried out as described in (Luecken and Theis, 2019), with thresholds for genes per cell and transcripts per cell set manually for each individual sample. The thresholds are summarized in supplemental table 2. In *S. purpuratus* samples, we were able to introduce another quality control feature and remove nuclei that had more than 5% of their transcripts originating from genes located on the mitochondrial scaffold. The *P. miniata* mitochondrial scaffold has yet to be identified, and therefore we could not introduce this metric to its quality control. For the previously published datasets from Foester et al. 2020, we used the published thresholds, with the addition of our 5% mitochondrial transcript limit.

### Creation of single-cell/nucleus atlas

Using Seurat V4.0.5 (Hao et al., 2021), each time point was individually normalized using the SCTransform algorithm (Hafemeister and Satija, 2019) while also regressing out the expression levels of rRNAs (as our library construction relied on polyA tagging). In our *S. purpuratus* samples, we also regressed out the percent mitochondrial transcripts. For the integration of individual samples, we used canonical correlation analysis (CCA) to identify up to 3,000 genes best suited to anchoring the datasets together using the SCT normalized values (Note: only 2,313 anchors were identified in our multi-species atlas). Following integration, Principal Component Analysis was run on the composite object and significant Principle Components (PCs) were identified by plotting the standard deviation of the first 50 PCs and identifying the PCs that describe the most variability of the datasets.

Our *P. miniata* nuclei atlas used 20 PCs, our *S. purpuratus* nuclear atlas used 10 PCs, the *S. purpuratus* whole-cell atlas used 10 PCs, and our multi-species atlas used 10 PCs. Using our selected PCs, we constructed k-nearest neighbor graphs (with k=20), performed Louvian Community Assignment (resolution set as 2 for *P. miniata* data, 1.5 for *S. purpuratus* nuclei data, 1.2 for *S. purpuratus* whole-cell data, and 1 for multi-species data), and performed Uniform Manifold Approximation and Projection (UMAP) reduction in order to generate clusters.

### Calculating marker genes

To annotate cluster identity, we identified genes that differentially marked clusters using the Wilcox rank-sum test. A gene is considered a marker gene for a cluster if it is expressed in at least 5% of the cells in that cluster and has at least a 0.25 log fold-change difference in average expression between the cells in that cluster and all other cells. Adjusted p-values were calculated using Bonferroni correction. We then used these marker genes to determine cluster cell-type labels by cross-referencing them with genes established in previous studies to have cell-type-specific expression.

### Ortholog identification and multi-species integration

1:1 orthologs were identified between *S. purpuratus* genome V 5.0 and *P. miniata* genome V3.0 and named according to orthologs identified in the human genome. For a full description of ortholog identification see Foley et al, 2021. To facilitate dataset integration, where applicable, the gene IDs in *P. miniata* datasets were converted to their orthologous *S. purpuratus* gene IDs. In total, 5790 of the 5935 1:1 orthologous genes identified between both species were expressed in at least 2 cells/nuclei in a dataset.

### Multi-species cluster composition and comparisons

To facilitate this comparison we constructed an *S. purpuratus* atlas at comparable time points to our *P. miniata* set (hatched and mesenchymal blastula stages) from a previously published dataset (Foster et al., 2020). CellRanger V 4.0.0 (Zheng et al., 2017) was used to align reads to a pre-mRNA index created using the *S. purpuratus* genome v5.0. After quality control (selection of transcript and gene count thresholds as with the *P. miniata* atlas, as well as the removal of cells with more than 5% of their transcripts originating from the mitochondrial scaffold), 7,370 cells were selected for analysis and clustering using Seurat V4.0.5 (Hao et al., 2021).

For each timepoint, reads were normalized using the SCTransform algorithm (Hafemeister and Satija, 2019), while regressing out mitochondrial (for *S. purpuratus* only) and rRNA transcripts. 2321 integration anchors were identified and used for dataset integration via CCA. UMAP dimensional reduction and Louvain community assignment resulted in 20 distinct clusters. To compare marker genes between different clusters, all calculated marker genes for a cluster of interest in their species-specific atlas were input into a Venn Diagram tool, housed by the Bioinformatics and Evolutionary Genomics group at the Ghent University (http://bioinformatics.psb.ugent.be/webtools/Venn/).

To calculate the significance of marker gene overlap between species, we calculated the representation factor of pairwise comparisons between calculated marker gene lists. It is calculated by dividing the number of overlapping genes by the expected number of overlapping genes for two independent sets subsampled from a larger finite set. Values over 1 indicate more overlap than expected by chance, while values under 1 indicate less overlap than expected. For marker comparisons between clusters in *P. miniata,* a total gene number of 27,818 was used. For cross-species analysis, this was calculated with respect to the total numbers of 1:1 orthologs. A calculator housed at http://nemates.org/MA/progs/representation.stats.html was used to conduct this analysis.

### Whole Mount In Situ Hybridization (WMISH)

The WMISH protocol was adapted from (Hinman et al., 2003b). Briefly, embryos were fixed in MOPS-4%PFA buffer and stored in 70% ethanol at -20°C. In situ primers for genes of interest were designed using Primer3 (Koressaar and Remm, 2007). Sequences were amplified from cDNA and DIG-labeled RNA probes complementary to the target mRNA (Roche). The primer sequences for synthesizing the probes are available in supplementary table 4. Three non-overlapping probes were created for *P. miniata’s VEGFR1L*.

Fixed embryos were brought to room temperature, placed into in situ buffer (0.1M MOPS, 0.5M NaCl, 0.1% Tween 20) washed twice, then transferred into hybridization buffer (0.1M MOPS, 0.5M NaCl, 1mg/mL BSA, 70% formamide, 0.1% Tween 20) and pre-hybridized overnight at 58°C. Following denaturation for 5 min at 65°C, DIG-labeled probes were added to the embryos at 0.2ng/µL. Samples were then incubated at 58°C for 4-7 days. Embryos were washed three times with a hybridization buffer over 4-6 hours, transferred to MAB (0.1M Malic Acid, 0.15M NaCl) with 0.1%Tween20 then washed four times with MAB + 0.1%Tween. Samples were blocked with MAB-2% Roche Block solution for 30 minutes at room temperature or overnight at 4°C. Anti-DIG AP FAB Antibody in MAB-2% Roche Block was added to the samples for a final concentration of 1:1000 and left to incubate for 2 hours at room temperature or overnight at 4°C.

Next, the samples were washed 4 times in MAB-0.1%Tween20 followed by three washes in AP buffer (0.1M NaCl, 0.1M Tris pH 9.5), with 15 minutes between washes. Samples were resuspended in color reaction solution (1X AP buffer, 50mM MgCl2, 0.2% Tween20, 0.175mg/µL 4-Nitro blue tetrazolium chloride, 0.35 mg/mL 5-Bromo-4-chloro-3-indolyl-phosphate), transferred to watch glasses, and kept in the dark at room temperature until the reaction was completed. When the color reaction concluded, embryos were washed three times in MAB-0.1%Tween20 and stored indefinitely in MAB.

Embryos were evaluated using a Leica MZ 95 dissection scope. For imaging, embryos were resuspended in MAB- 50% glycerol, and photographs were taken using a Leica DMI 4000 B inverted scope and Leica Application Suite X V3.6.20104.

### Image Creation and Processing

Gimp was used to process microscopy images (The GIMP Development Team, 2021). BioRender.com was used to create schematics in Figure 1.A and Figure 8.A. Inkscape was used to compile and format the final images (Inkscape Project, 2021).

---

**Supplemental Figure 1.**
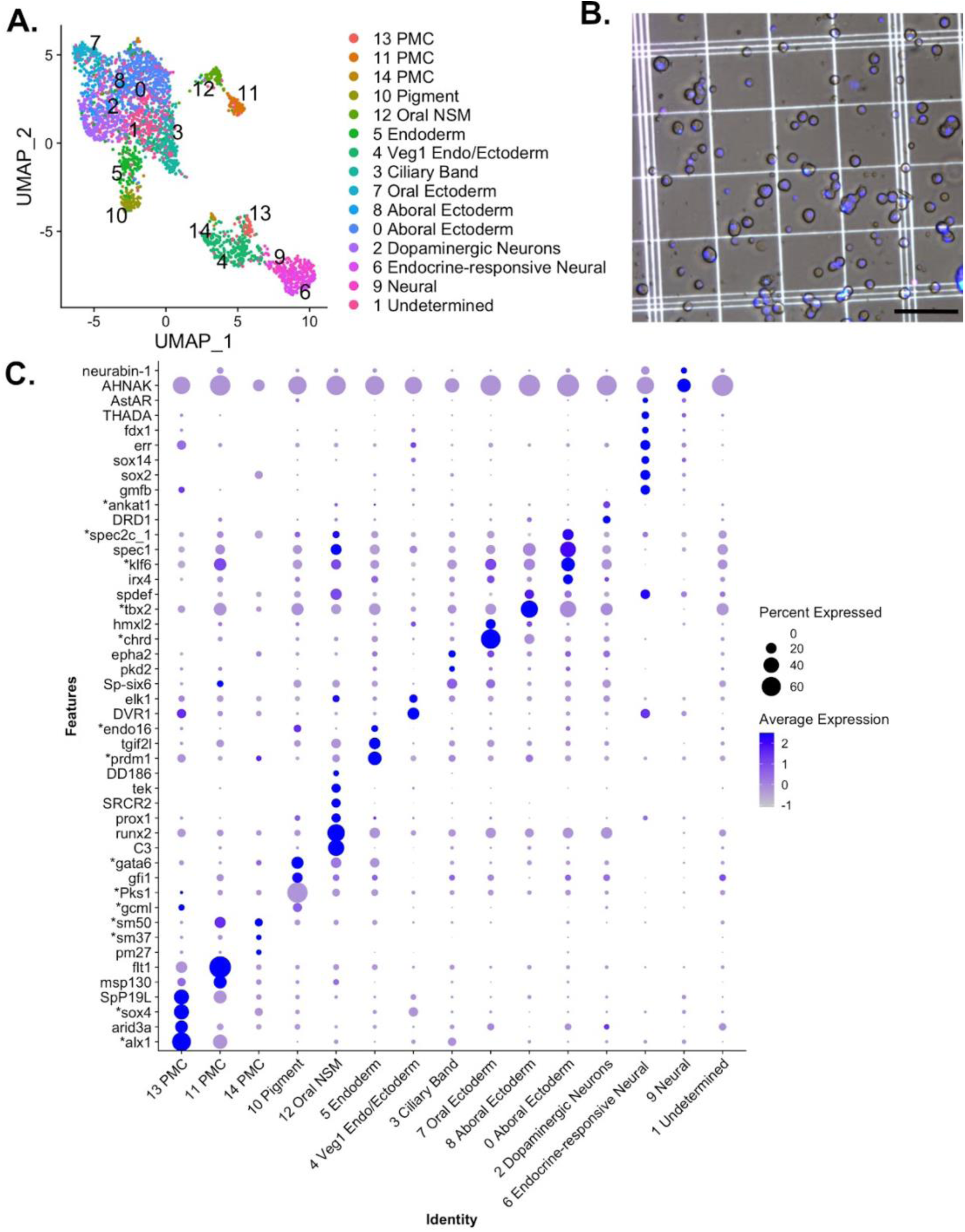
Single nucleus atlas of early development in S. purpuratus. (A) A UMAP reduction of the combined *S. purpuratus* single nucleus datasets results in 15 clusters. (B) An example of a nuclear isolate. DNA is stained with Hoescht dye. The scale bar represents 250 μm. (C) A dot plot highlighting genes used to annotate clusters. Along the X-axis are the cluster names. Along the Y-axis are the genes. The size of a dot is proportional to the percentage of nuclei in that cluster expressing a given gene, while the shade of the dot correlates with the expression level. Genes that were also used as markers in Foster et al 2020 are indicated with an asterisk.

**Supplemental Figure 2.**
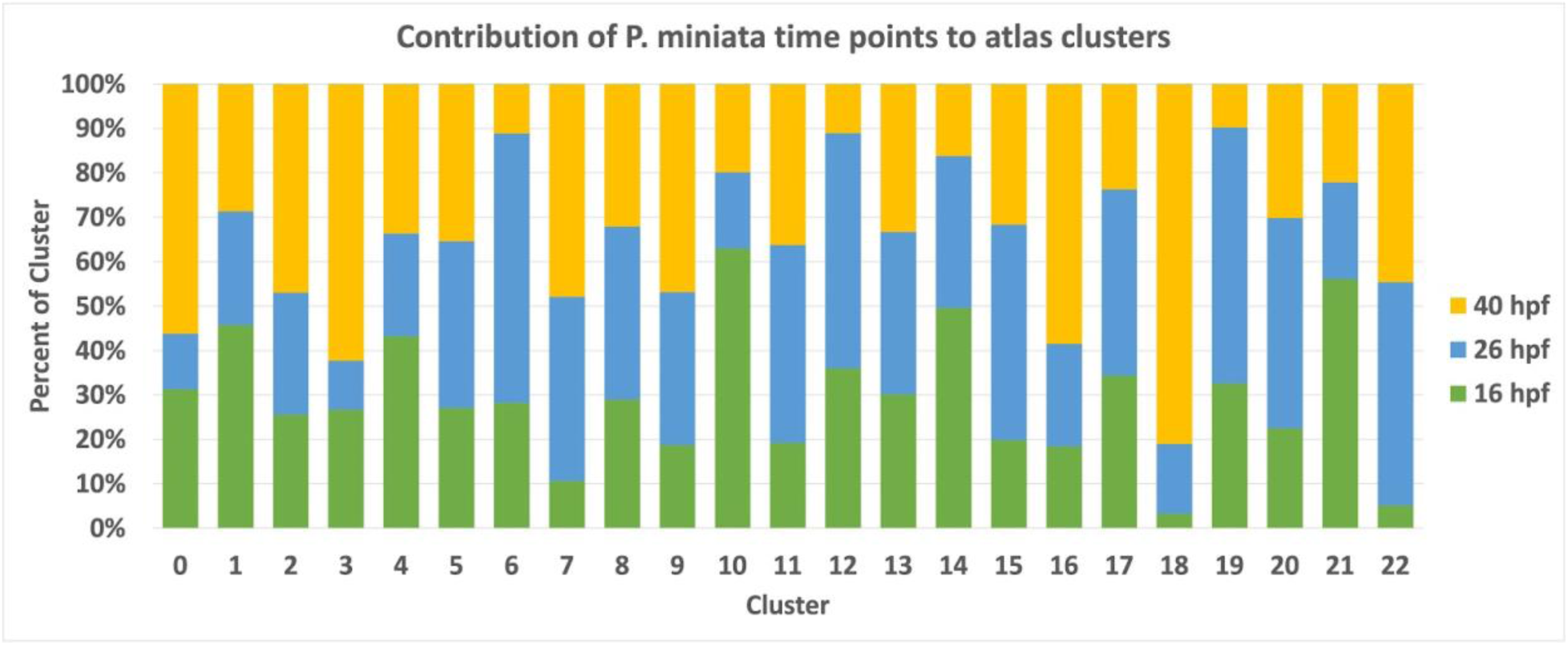
Contribution of *P. miniata* time points to atlas clusters. A stacked bar graph displaying the breakdown of each cluster in the *P.miniata* developmental atlas by time point of sampling, normalized by sample size.

**Supplemental Figure 3:**
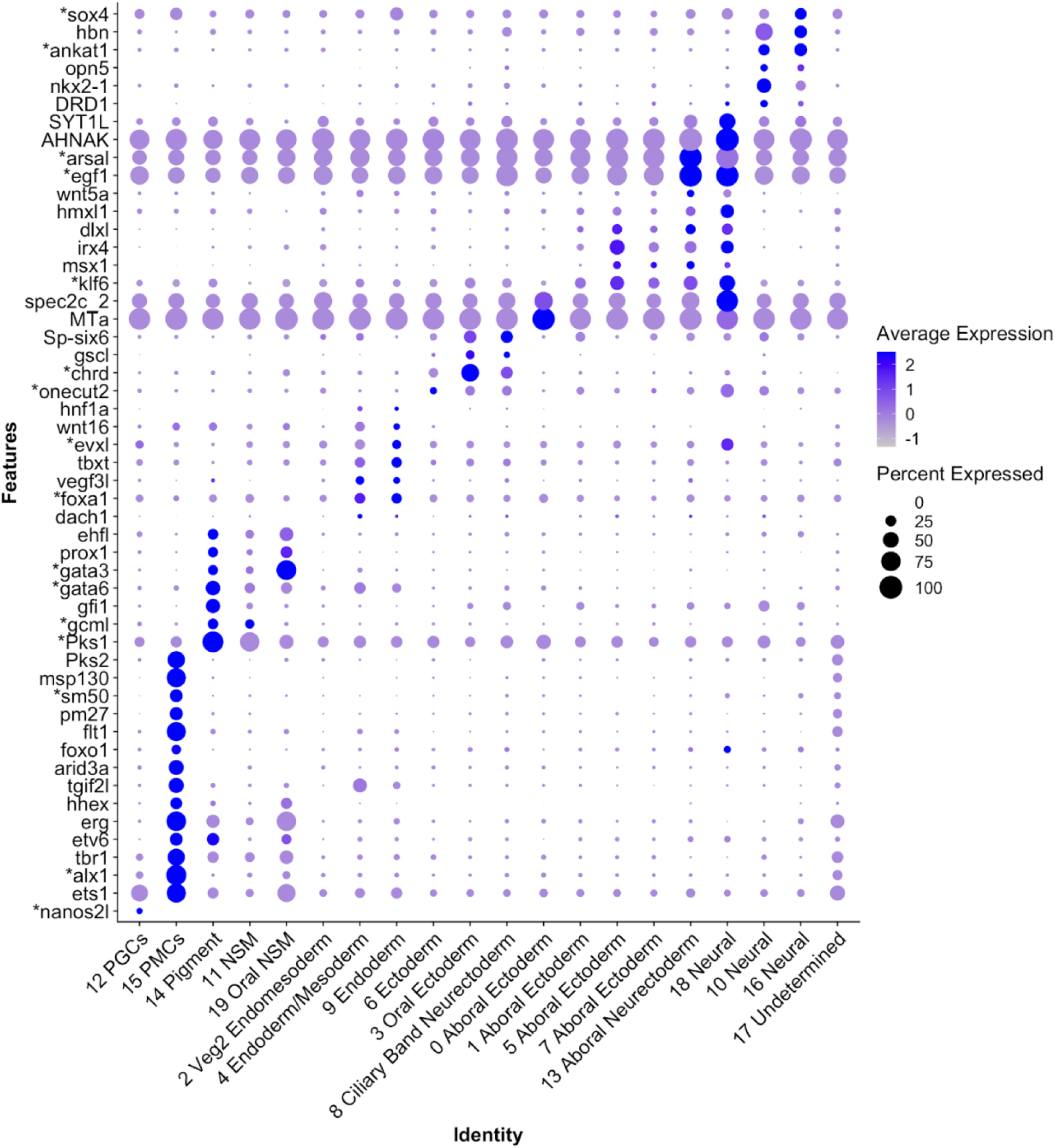
Differential expression of marker genes used to annotate *S. purpuratus* atlas. A dot plot highlighting marker genes used to determine cluster identity. The X-axis lists the cluster names, while the Y lists gene names. An asterisk indicates it was also defined as a marker in Foster et al 2020. Circle size corresponds to the number of cells in the cluster expressing the gene of interest, while shade correlates with the level of expression.

