## Supplemental Text for "New hypotheses of cell type diversity and novelty from comparative single cell and nuclei transcriptomics in echinoderms"

### Supplemental Section 1: A single nucleus atlas of early development in *S. purpuratus*

As a proof of concept, we performed single-nucleus RNA-seq on *S. purpuratus* embryos to see if we could create biologically meaningful clusters that correspond to known cell types. We sample 4 time points spanning the blastula through the mid-gastrula stage (6hpf, 15hpf, 23hpf, and 33hpf)

Of particular note is that our final clustering resulted in 3 PMC clusters, whereas previous whole-cell work has only yielded 1 PMC cluster without conducting any sub clustering analysis [(Paganos et al. 2021; Foster et al. 2020)](https://sciwheel.com/work/citation?ids=12089028,12128570&pre=&pre=&suf=&suf=&sa=0,0). Cluster 13 expresses aristaless-like homeobox (alx1), AT-rich interactive domain-containing protein 3A (*arid3a*, previously called dead ringer homolog), and SRY-box transcription factor 4 *(sox4)*, genes in the most upstream, an early-acting portion of the PMC GRN. Cluster 11 is marked by the expression of vascular endothelial growth factor receptor 1 *(flt1,* also known as *VEGFR1*), a gene previously implicated in activating biomineralizing effector genes, as well as mesenchyme-specific cell surface glycoprotein (*msp130*), a gene present in the biomineralization module of the PMC GRN. Cluster 14 is characterized by the expression of other genes involved in the process of biomineralization, including spicule matrix protein SM37 (*sm37*), and spicule matrix protein SM37 (*sm37*). The distinction of our PMC clusters, consistent with the order of GRN deployment suggests that nuclei give us an advantage for detecting subpopulations of this morphologically complex cell type.

Pigment cells were marked b the expression of Zic family member 1 (*zic1*), glial cells missing transcription factor-like protein (*gcml*), and GATA binding protein 6 (*gata6*). It also shows broad expression of probable polyketide synthase 1 (*Pks1*).

The Oral NSM is characterized by the expression of known blastocoel cell markers like RUNX family transcription factor 2 (*runx2*), and prospero homeobox 1(*prox1*). This identity is further supported by the expression of genes related to immune function, including complement C3 (*C3*), scavenger receptor cysteine-rich protein variant 2 (*SRCR2*), TEK receptor tyrosine kinase (*tek*) and DD186 protein, upregulated upon bacterial challenge (*DD186*).

Cluster 5 was annotated as endoderm because it is clearly distinguished by the expression of PR/SET domain 1 (*prdm1*), homeobox protein TGFB induced factor homeobox 2-like (*tgif2l*), and calcium-binding protein (*endo16*)

Cluster 4 corresponds to the ectodermal and endodermal veg1 region of the embryo is marked by the expression of protein DVR-1 homolog (*DVR1*) and transcription factor ETS transcription factor ELK 1 (*elk1*).

Cluster 3 Ciliary band is characterized by the expression of homeobox protein SIX homeobox 6 (*Sp-six6*), polycystin 2, transient receptor potential cation channel (*pkd2*), and EPH receptor A2 (*epha2*).

The oral ectoderm is marked by the strong expression of chordin (*chrd*) as well as H6 family homeobox-like 2 (*hmxl2*)

Clusters 8 and 0 are annotated as aboral ectoderm and are characterized by the expression of T-box transcription factor 2 (*tbx2*), SAM pointed domain-containing ETS transcription factor (*spde*f, formerly known as *Ets4*), iroquois homeobox 4 (*irx4*), Kruppel like factor 6 (*klf6*), spec 1a protein (*spec1a*), and spec 1a protein (*spec2c_1*).

Three neural clusters were identified in our single nucleus atlas. . Cluster 6 was annotated as endocrine-responsive neural because this cluster expresses a variety of transcription factors involved in neural differentiation, including SRY-box transcription factor 2 (*sox2*), SRY-box transcription factor 14 (*sox14*), H2.0-like homeobox protein (*hlx*), and glial cells missing transcription factor-like (*gcml*). This cluster also expresses genes linked to hormone signaling, estrogen related receptor (*err*), allatostatin-A receptor (*AstAR*), and membrane-associated progesterone receptor component 1-like (*PGRMC1L*).

Cluster 2 was identified as dopaminergic neurons based on the marker gene activity of D(1) dopamine receptor (*DRD1*) and ankyrin containing gene specific for apical tuft 1 (*ankat1*).

Cluster 9 is also annotated as neural-based on the expression of neuroblast differentiation-associated protein AHNAK (*AHNAK)* and *neurabin-1* and is specific neuronal lineage cannot be determined based on the marker genes expressed at these stages of development.

Only cluster 1 was not able to be annotated, as it expressed no previously-characterized cell type markers.

The ability to resolve these diverse cell stages at such an early point in development, despite the small size of the dataset, demonstrates the strengths of a single nucleus approach to developmental atlas creation.

### Supplemental Section 2: Detailed cluster annotation for P. miniata sn-RNA-seq

**Cluster 0** had only two genes in its marker set (nuclear receptor subfamily 0 group B member 1-like and filamin-B) and we therefore overall had low confidence in our ability to annotate this cluster. We characterized this cell cluster as **mesoderm-like** based on absence of markers. That is, key endoderm (*gata6, arid3a*, T-box transcription factor 2 (*tbx2*)) and ectoderm (ets variant transcription factor 6 (*etv*), *PM-onecut2* ) underexpressed in cluster 0 compared to other clusters.

**Cluster 1** we annotaed as **ectoderm** based on the presence of the one cut domain family member 2-like (*PM-onecut2*) , goosecoid homeobox-like (*gscl*), and forkhead box j1 (*foxj1*), all of which are known to be expressed in *P. miniata* ectoderm [(Yankura et al. 2010)](https://sciwheel.com/work/citation?ids=402624&pre=&suf=&sa=0) .

Cluster 2 was annotated as ectoderm, based on presence of the marker genes, *Pm-onecut2* and orthodenticle homeobox 2 (*otx2*). *Foxj1* and *Gsc* were also expressed. This cluster was therefore similar to cluster 1, with significant expression of *otx2*, and higher levels of expression of *onecut2* making it distinct from cluster 1

Prdm1 (formely known as *Blimp1/krox*) which is expressed in the veg1 ectoderm/endoderm boundary is marker gene of **cluster 3**. We also detected low levels of *gata6, tgif2l, foxa1,* and *foxn3* which are all expressed in the endodem. We therefore annotated cluster 3 as **veg1 ectoderm/endoderm cells**.

**Cluster 4** was annotated as **post-oral ciliary band ectoderm**. A marker of this cluster is *Foxj* which is broadly expressed in the ectoderm. *Nk1-2l* and *gscl* also show low levels of expression in this cluster and are specifically expressed in the post-oral cilairy band. Additionally, multiple cytoskeletal and motor protein associated genes appear in this clusters marker gene set, including stabilizer of axonemal microtubules 1-like (*saxo1*), stabilizer of axonemal microtubules 2 (*saxo2*), and dynein heavy chain 2, axonemal-like (*DNAH2L*),and tectin b1.

The ventral ectoderm expressed gene *tbx2* is a strong marker gene in cluster 5 We also noted expression of mothers against decapentaplegic homolog 6-like (*smad5l*) as known marker of ventral ectoderm in sea urchins. It is also markers by genes relating to ciliary band function like cilia and flagella associated protein 410 (*CFAP410*)and dynein heavy chain 2, axonemal-like (*DNAH2*). This cluster was here annotated as ventral ectoderm

**Cluster 6 was annotated as muscle/coelom.** It was marked by the expression of myosin light chain kinase smooth muscle-like (*MLCKL*). Secreted frizzled-related protein 5-like (SFRP5L), Zic family member 1 (*zic1)*, homeobox protein *SIX6-like (Six6L)*, and *ets1* are also lowly expressed in this cluster. *Frzl-5* is expressed in the anterior coelom, the source of muscle cells.

**Cluster 7** was marked by the expression of has chordin-like (chrdl), *gscl, arid3a*, and transcription factor AP-2 alpha *(tfap2a*). In situ hybridization (Fig 7.D.vi) has shown *tfap2a* localizes to the oral ectoderm. We annotated this cluster as **oral ectoderm**.

**Cluster 8** shows expression of multiple veg1 ectoderm genes, including Wnt family member 3 (*wnt3*) and transcription factor HES-4 (*HES4*). The presence of NK1 transcription factor-related protein 2-like (nk1-2l) is also detected. The gene lim homeobox 1 (*lhx1*) is highly expressed in its marker gene set. *lhx1* is also expressed in the ectoderm of *S. purpuratus*. Based on this, we determined cluster 8 is **veg1 ectoderm cells.**

Like cluster 7, **cluster 9** has *tfap2a* and *arid3a* in its marker gene set. In comparison to cluster 7, cluster 9 includes *epha2* as a maker gene, which is expressed in ciliary band [(Krupke and Burke 2014)](https://sciwheel.com/work/citation?ids=260725&pre=&suf=&sa=0). Therefore we also annotate **cluster 9 as oral ectoderm.**

**Cluster 11**  was annotated as **veg1 ectoderm/endoderm**. *Hbox 7, gata6, Wnt3*, and *prdm1* are in this clusters marker gene set, all of which are in the *P. miniata* veg1 endoderm/ectoderm GRN. Cluster 11 was distinct from clusters 3 and 8 based on the maker gene *wnt1* and strong expression of *hbox7*

**Cluster 12** was not marked by the expression of any genes with pre-existing literature in the sea star and thus we were unable to assign it an identity. It remains **undetermined**. Its top 3 marker genes tha have been named are organic cation transporter protein-like (*Orctl)*,probable beta-D-xylosidase 6 (*BXL6*), and poly [ADP-ribose] polymerase tankyrase-like (*TNKS2L*).

**Cluster 13** has *gata6, prdm1, foxn3,* and *tgif2l*  in its marker gene set, all of which are part of the P. miniata endoderm GRN. Compared to cluster 3, this higher expression of *gata6* and presence of *wnt3.* *Wnt3* is strongly expressed in endoderm [(McCauley et al. 2013)](https://sciwheel.com/work/citation?ids=1308627&pre=&suf=&sa=0). Thus, this cluster is defined as **endoderm.**

**Cluster 15** has hedgehog family (*hh*) and fibroblast growth factor receptor (*fgfrl1*) in its marker gene set, which are both expressed in the foregut . Smoothelin-like (*SMTN*L) and forkead box A1 (*foxa*) are also expressed in this cluster. Therefore, we define cluster 15 as the **foregut**.

**Cluster 19** is marked by the expression of s*ox2, sox14*, and several cell cycle regulators, including geminin-like (*GMNNL*),PCNA-associated factor-like (*PCLAFL*), chromatin licensing and DNA replication factor 1 (*cdt1*), cyclin A2 (*ccna2*), and cyclin =B2 (*ccnb2*). Therefore, we define cluster 19 as **dividing ectoderm.**

**Cluster 20** is marked by the expression of *foxj1*, several genes relating to cilia and flagella activity. These include members of the tektin family (tektin 2 (*tekt2*), tektin 3 (*tekt3*), and tektin 4 (*tekt4*)), and radial spoke head component 1 (*rsph1*). We, therefore, annotate this cluster as broad **ciliary band.**
